# Tau-mediated Disruption of the Spliceosome Triggers Cryptic RNA-splicing and Neurodegeneration in Alzheimer’s Disease

**DOI:** 10.1101/514927

**Authors:** Yi-Chen Hsieh, Caiwei Guo, Hari K. Yalamanchili, Measho Abreha, Rami Al-Ouran, Yarong Li, Eric B. Dammer, James J. Lah, Allan I. Levey, David A. Bennett, Philip L. De Jager, Nicholas T. Seyfried, Zhandong Liu, Joshua M. Shulman

## Abstract

In Alzheimer’s disease (AD), spliceosomal proteins with critical roles in RNA processing aberrantly aggregate and mislocalize to Tau neurofibrillary tangles. We test the hypothesis that Tau-spliceosome interactions disrupt pre-mRNA splicing in AD. In human postmortem brain with AD pathology, Tau coimmunoprecipitates with spliceosomal core components. In *Drosophila* models, pan-neuronal Tau expression triggers reductions in core and U1-specific spliceosomal proteins, and genetic disruption of these factors, including SmB, U1-70K, and U1A, enhances Tau-mediated neurodegeneration. We further show that loss-of-function in *SmB*, encoding a core spliceosomal protein, causes decreased survival, progressive locomotor impairment, and neuronal loss, independent of Tau toxicity. Lastly, RNA-sequencing reveals a similar profile of mRNA splicing errors in *SmB* mutant and Tau transgenic flies, including intron retention and non-annotated cryptic splice junctions. In human brains, we confirm cryptic splicing errors in association with neurofibrillary tangle pathologic burden. Our results implicate spliceosome disruption and perturbations of the neuronal transcriptome in Tau-mediated neurodegeneration in AD.

## INTRODUCTION

In eukaryotes, precursor messenger RNA (pre-mRNA) splicing removes introns and generates mature mRNA transcripts, subserving a critical role in the regulation of gene expression. Splicing contributes to neuronal transcriptional diversity and function, and disruption of splicing mechanisms causes neurologic disease (Cooper et al., 2009; Tollervey et al., 2011). For example, spinal muscular atrophy is caused by mutations in the *survival motor neuron* (*SMN*) gene, which initiates assembly of the spliceosome, the molecular machine responsible for pre-mRNA splicing (Lefebvre et al., 1995; Lorson et al., 1999). Mutation of other RNA-binding proteins implicated in splicing, including the TAR DNA-binding protein 43 (TDP-43) and fused in sarcoma (FUS), are associated with frontotemporal dementia and amyotrophic lateral sclerosis (FTD-ALS) (Neumann et al., 2006; Sreedharan et al., 2008; Vance et al., 2009). In mouse experimental models, loss-of-function in spliceosomal components, is also associated with neurodegenerative phenotypes (Jia et al., 2012; Ling et al., 2015; Polymenidou et al., 2011; Tan et al., 2016; Zhang et al., 2008).

The major spliceosome is comprised of 5 small nuclear ribonucleoprotein (snRNP) complexes (U1, U2, U4, U5, and U6), each including a small nuclear RNA (snRNA), seven Sm proteins (or Lsm proteins in U6), and specific factors, such as U1-70K, U1A, and U1C for the U1 snRNP (Will and Luhrmann, 2011; Matera and Wang, 2014). Assembly begins with the formation of the core snRNA-Sm protein complex in the cytoplasm, followed by nuclear import and subsequent incorporation of specific proteins to generate the mature snRNP. Pre-mRNA splicing is initiated by recognition of 5’ splice sites by the U1 snRNP, followed by dynamic interactions with other snRNP complexes. Disruption of spliceosomal factors, either in cell culture or mouse genetic models induces widespread mRNA splicing errors, including intron retention and cryptic junctions—consisting of non-conserved, non-annotated splice junctions (Humphrey et al., 2017; Jia et al., 2012; Kaida et al., 2010; Ling et al., 2015; Polymenidou et al., 2011; Tan et al., 2016; Zhang et al., 2008). Cryptic splicing has also been documented in human postmortem brain from individuals with TDP-43 mutations (Ling et al., 2015). In fact, emerging evidence suggests that disrupted assembly of RNA-binding protein complexes, such as the spliceosome, may promote FTD-ALS pathogenesis (Ito et al., 2017; Lee et al., 2016; Yin et al., 2017).

Tau protein, encoded by the *microtubule associated protein tau* gene (*MAPT*), aggregates to form neurofibrillary tangles (NFTs), characteristic of Alzheimer’s disease (AD) and other tauopathies. Neurofibrillary tangle pathologic burden is strongly correlated with cognitive decline in AD (Arriagada et al., 1992; Braak and Braak, 1991; Gómez-Isla et al., 1997), and soluble, oligomeric forms of Tau are also implicated in synaptic dysfunction and neuronal death (Cowan and Mudher, 2013; Spires-Jones and Hyman, 2014). In human AD postmortem brain tissue, multiple core and specific components of the U1 snRNP co-aggregate with Tau in neurofibrillary tangles (Bai et al., 2013; Hales et al., 2014), and similar findings have been reported in *MAPT* transgenic mice (Maziuk et al., 2018; Vanderweyde et al., 2016) and *in vitro* (Bishof et al., 2018). Consistent with these observations, evidence of altered splicing in AD has also recently emerged (Bai et al., 2013; Raj et al., 2018). Independently, in a screen of candidate genes from AD-associated human genomic loci, we discovered that *SmB*, the fly ortholog of human *SNRPN*/*SmN*, modulated Tau-mediated neurotoxicity (Shulman et al., 2014). Here, we couple studies in human autopsy cohorts and *Drosophila* models to further investigate the hypothesis that Tau-spliceosome interactions lead to splicing errors and ultimately, neurodegeneration in AD.

## RESULTS

### Tau associates with numerous snRNP core components in human brains with AD pathology

We previously showed that multiple core and U1-specific components of the spliceosome are enriched in insoluble protein fractions and closely associate with neurofibrillary tangles in AD postmortem brain tissue (Bai et al., 2013; Bishof et al., 2018; Hales et al., 2014). To further explore the potential for interactions with soluble, oligomeric forms of Tau that most likely mediate toxicity, we performed immunoaffinity-purification coupled to mass spectrometry. A Tau monoclonal antibody (Tau5) was used for immunoprecipitation in human brain lysates from AD and non-demented control subjects (n = 4 each, Table S1). As a negative control, we performed IP with a non-specific IgG from pooled control and AD inputs. Tau immunoprecipitation was confirmed by western blot analysis (Figure 1A). Next, samples were on-bead digested and peptides analyzed by liquid chromatography-tandem mass spectrometry (LC-MS/MS) using label-free quantitation (LFQ). Our analysis identified 1,065 proteins across all samples. Differential enrichment analysis of Tau-interacting partners identified 513 proteins enriched in AD versus control brains (*p* < 0.05, minimum 1.5-fold change, Table S2), highlighting proteins with significantly altered interactions in the context of AD pathology (Figure 1B). Among those proteins characterized by increased affinity for Tau in AD are numerous ribonucleoproteins (p = 1.3×10^−14^) based on DAVID gene ontology enrichment analysis, with roles in mRNA processing, including splicing and/or translation (Figure 1B, Figure S1, and Tables S2 & S3). Among these, 10 spliceosome proteins, including SNRNP70 (U1-70K), SNRPD2 (SmD2), SNRPD3 (SmD3), SNRPN (SmN), and SNRPA (U1A), each exhibited more than 3-fold increased affinity to Tau in AD brains versus control (Figure 1C). These data suggest that in AD, soluble forms of Tau may associate with spliceosome components, possibly preceding the formation of NFTs.

**Figure 1.**
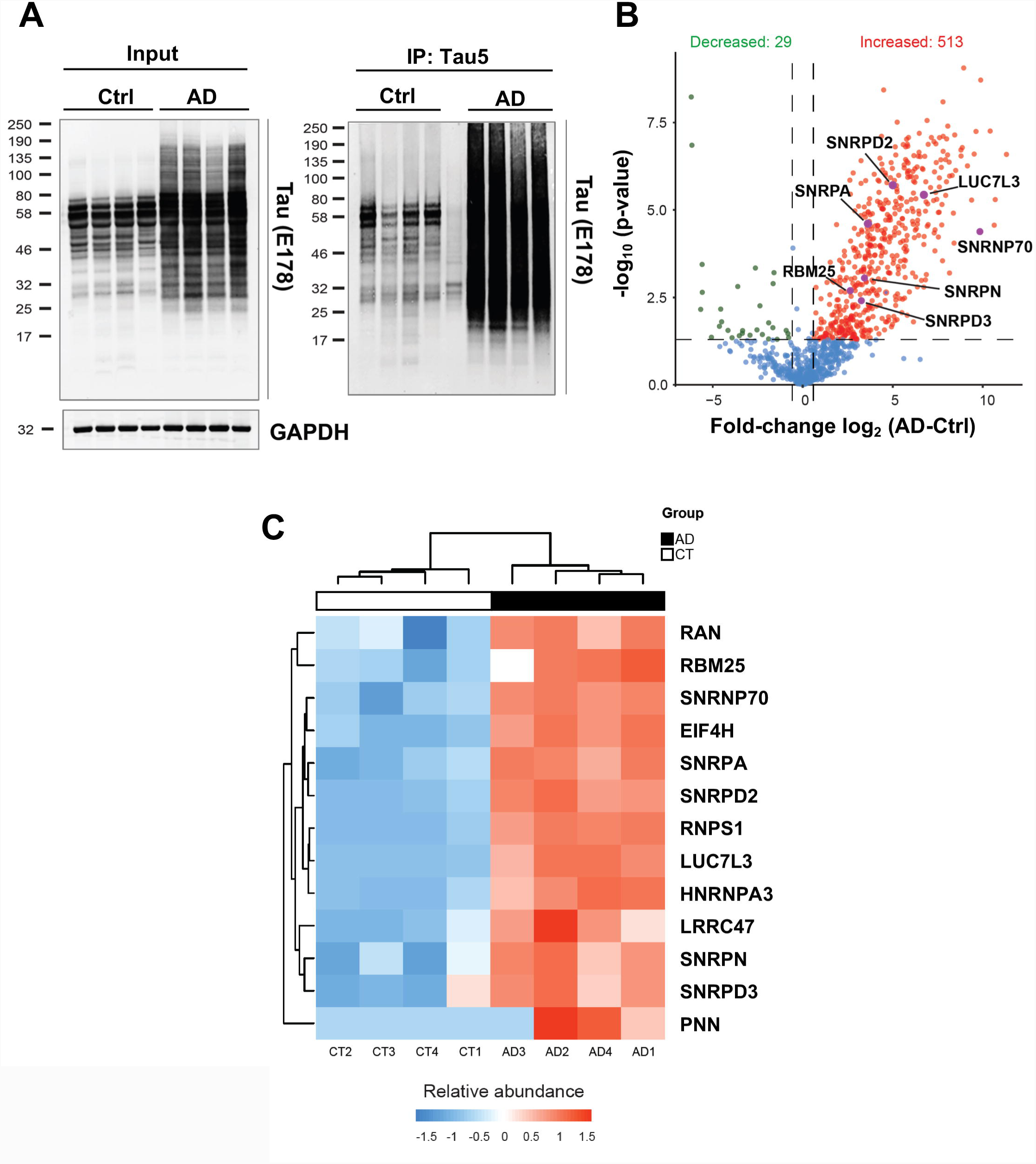
Tau associates with spliceosome components in human AD postmortem brain. (A)Tau was immunoprecipitated (IP) using the Tau5 monoclonal antibody from either healthy control (n = 4) or AD postmortem (n = 4) brain homogenates. Western blots were probed with the polyclonal antibody, Tau-E178, demonstrating accumulation of Tau protein in AD, including higher molecular weight species. See also Table S1. **(B)** Liquid chromatography coupled to tandem mass spectrometry was performed, identifying 1,065 proteins co-immunoprecipitating with Tau. Volcano plots highlight Tau-associated proteins exhibiting differential interactions in AD versus control brains, including those with 1.5-fold or greater increased (Red, n = 513) or decreased (Green, n = 29) affinity. Among those proteins with increased affinity for Tau in AD are numerous ribonucleoproteins (p = 1.3×10^−14^) based on DAVID gene ontology enrichment analysis. Multiple spliceosome components are highlighted. See also Tables S2 & S3. **(C)** Heatmap highlighting representative Tau-associated ribonucleoproteins, including multiple spliceosome components, with increased (Red) interactions in AD cases (AD, Black) versus controls (CT, White). The 13 proteins depicted here comprise a tight protein-protein interaction subnetwork based on STRING analysis (Figure S1). All proteins are differentially associated with Tau (Fold-change > 1.5, p < 0.05). For statistical analysis, unpaired t-test was performed. For statistical analyses (**C** & **E)**, one-way-ANOVA was performed followed by Tukey’s test for post hoc comparisons. **, p < 0.01; ***, p < 0.001. All error bars denote mean ± SEM.

### Tau genetically interacts with multiple core spliceosomal components in ***Drosophila***

Expression of human *MAPT* in *Drosophila* is neurotoxic, including both wild-type *Tau* (*Tau*^*WT*^) or mutant forms associated with familial frontotemporal dementia. We initially selected a mutant *Tau*^*V*^*337*^*M*^ transgenic fly strain which is amenable to sensitive and robust detection of genetic modifiers (Shulman and Feany, 2003; Shulman et al., 2011, 2014). Using the *GMR*-*GAL4* driver, we directed expression of Tau to the *Drosophila* retina, along with RNA-interference (RNAi) transgenes targeting 10 distinct U1 snRNP components, including the Sm core (SmB, SmD1, SmD2, SmD3, SmE, SmF, and SmG) and U1-specific factors (U1-70K, U1C, and U1A). We required consistent interactions with at least 2 independent lines to minimize off-target effects and excluded RNAi strains with evidence of significant retinal toxicity in the absence of Tau (Table S4). These experiments confirmed our prior results with *SmB* (Shulman et al., 2014) and additionally revealed that knockdown of fly homologs of *U1-70K, U1C, SmD2*, and *SmE* similarly enhance Tau-induced retinal degeneration, causing reduced eye size and increased, roughened appearance (Figure 2A). We next employed a complementary assay in which *Tau* expression is restricted to adult photoreceptors, using the *Rhodopsin 1* (*Rh1*)-GAL4 driver, causing an age-dependent, progressive loss of the light-induced depolarization response, but preserved retinal morphology (Chouhan et al., 2016). Based on electroretinogram (ERG) assays, we confirmed that RNAi-knockdown of U1 snRNP components showed consistent enhancement of the *Rh1>Tau*^*WT*^ functional degenerative phenotype (Figure S2B). To further examine for dose-sensitive genetic interactions, we also tested available mutant alleles, including available null alleles for fly *U1-70K* and *snf* (ortholog of *U1A*) (Flickinger and Salz, 1994; Salz et al., 2004) or a newly generated *SmB* hypomorphic allele (*SmB*^*MG*^, see below). The Tau ERG phenotype was dominantly enhanced in either an *SmB*^*MG*/+^ (Figure 2B-C) or *snf*^+/-^ heterozygous genetic background (Figure S2B) but not in *U1-70K*^+/-^ (Figure S2C), whereas control heterozygous animals had normal ERGs in the absence of Tau.

**Figure 2.**
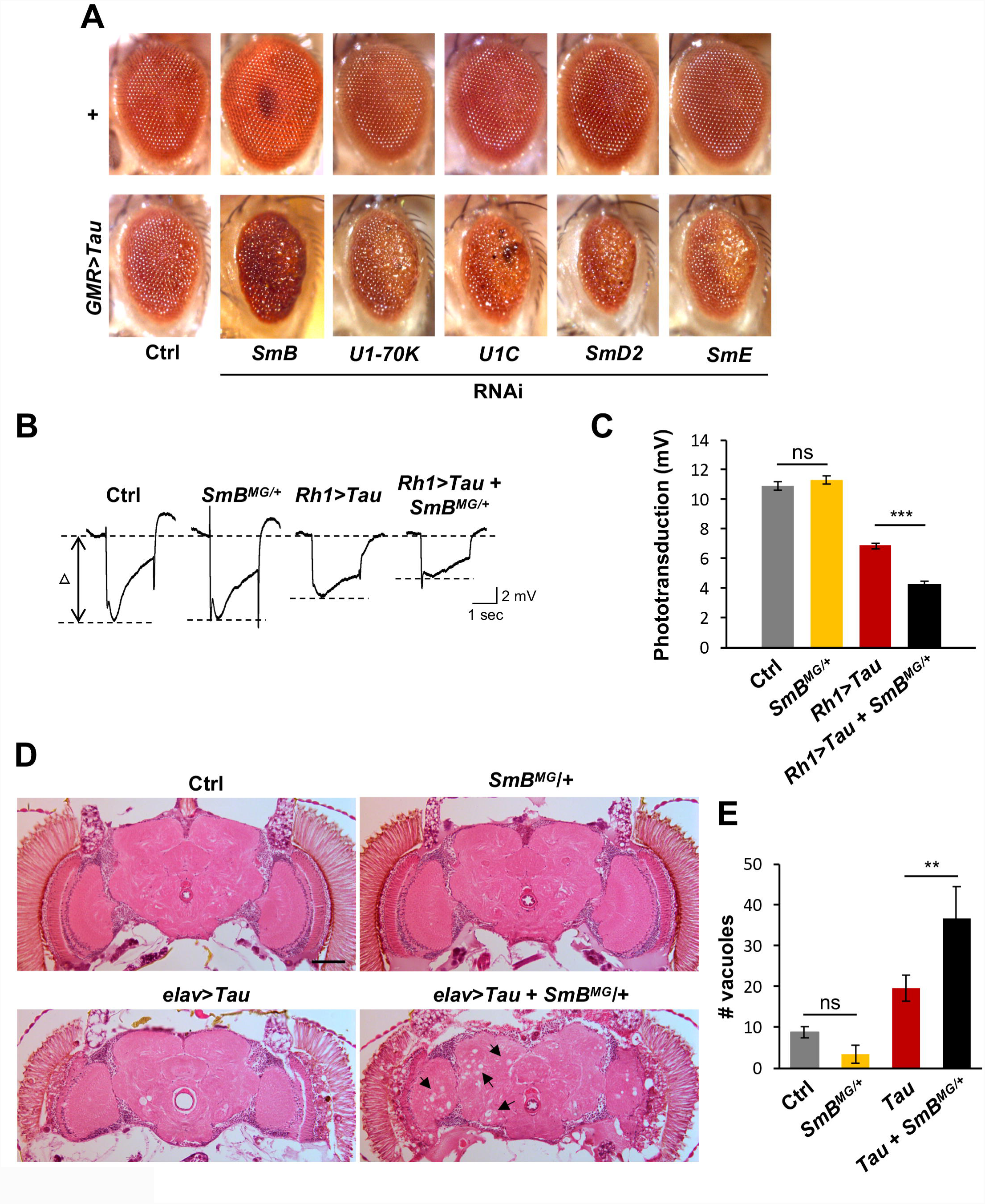
Genetic interactions between Tau and numerous core spliceosomal factors. (**A**) RNA-interference (RNAi) targeting multiple core and U1-specific spliceosome factors enhances Tau-mediated retinal toxicity. Expression of both Tau and RNAi were directed to the retina using the eye-specific driver, *GMR*-*GAL4*. Compared with control animals (Top Left: *GMR-GAL4*/+), expression of human Tau (Bottom Left: *UAS-Tau*^*V*^*337*^*M*^/+; *GMR*-*GAL4*/+) causes reduced eye size and roughened appearance. RNAi targeting multiple spliceosome components enhanced Tau retinal toxicity, exacerbating the rough eye phenotype (Bottom row: *UAS-Tau*^*V*^*337*^*M*^/+; *GMR*-*GAL4*/*UAS*-*RNAi* or *UAS- Tau*^*V*^*337*^*M*^/+; *GMR*-*GAL4*/*+; UAS-RNAi*/*+*), including *SmB*^*HM05097*^, *U1-70K*^*v23150*^, *U1C*^*v22132*^, *SmD2*^*v31947*^, and *SmE*^*v23570*^. RNAi lines were not associated with any significant toxicity when expressed independently of Tau (Top row: *GMR*-*GAL4*/*UAS-RNAi* or *GMR*-*GAL4/+; UAS-RNAi/+*). The following RNAi lines showed consistent interactions, providing further independent confirmation: *SmB*^*v110713*^, *U1-70K*^*HMS00274*^, *U1C*^*v22133*^, *U1C*^*HMS00137*^, *SmD2*^*v31946*^, *SmD2*^*HMC03839*^, and *SmE*^*HMS00074*^. See also Table S4. (**B**) *SmB* exhibits dose-sensitive enhancement of Tau-induced retinal dysfunction. Light induced depolarization of retinal photoreceptors was recorded using electroretinograms (ERGs) in 5-day-old animals. Compared to controls (*Rh1-GAL4*/+ and *Rh1-GAL4*/*SmB*^*MG*^), Tau expression in the adult retina, via the *Rh1-GAL4* driver, causes progressive loss of ERG amplitude (*Rh1-GAL4*/*+*; *UAS-Tau*^*WT*^/*+*), and this phenotype is enhanced in flies heterozygous for the *SmB*^*MG*^ hypomorphic allele (*Rh1-GAL4*/*SmB*^*MG*^; *UAS-Tau*^*WT*^/+). (**C**) Quantification of experimental data in (**B**) based on examination of 10 animals per genotype. See also Figure S2. (**D**) *SmB* exhibits dose-sensitive enhancement of Tau-induced neurodegeneration in the adult brain. Tau was expressed pan-neuronally using the *elav-GAL4* driver. Frontal sections were prepared from 10-day-old animals and stained with hematoxylin and eosin to reveal neurodegenerative changes. Compared with controls (Top Left: *elav-GAL4*/+ and Top Right: *elav-GAL4*/+; *SmB*^*MG*^/+), pan-neuronal expression of Tau (Bottom Left: *elav-GAL4*/+; +/+; *UAS-Tau*^*R406W*^/+) causes progressive neuropil vacuolization (arrows), and this phenotype is enhanced in flies heterozygous for the *SmB*^*MG*^ hypomorphic allele (Bottom Right: *elav-GAL4*/+; *SmB*^*MG*^/+; *UAS-Tau*^*R406W*^/+). Scale bar: 50 μm. (**E**) Quantification of vacuole formation in **C**, based on the examination of central brain sections from at least 8 brains per group (same genotypes denoted in **D**).

Next, we expressed Tau pan-neuronally using the *elav*-*GAL4* driver line, which causes age-dependent neuronal loss and associated histologic changes in the adult brain (Wittmann et al., 2001). RNAi knockdown of U1 snRNP genes using *elav-GAL4* resulted in embryonic lethality in most cases, so we again took advantage of the available mutant alleles to examine for dominant genetic interactions. In 10-day-old animals, *elav>Tau*^*R406W*^ causes a mild degree of neurodegenerative changes, based on the accumulation of vacuoles on hematoxylin and eosin stained, paraffin brain sections (Figure 2D). By contrast, Tau-induced neurodegeneration was dominantly enhanced in either a *SmB*^*MG*/+^ (Figure 2D-E) or *U1-70K*^+/-^ (Figure S2D-E) heterozygous genetic background, but not in *snf*^+/-^. We did not detect evidence of neurodegeneration in heterozygous *SmB* or *U1-70K* control flies independent of Tau (Figure 2D-E and Figure S2E). In sum, based on multiple independent assays, our data suggest that genetic manipulation of U1 snRNP components can enhance Tau-induced neurodegenerative phenotypes in *Drosophila*.

### Tau-induced spliceosome disruption in ***Drosophila***

Given the observed genetic interactions, we next examined the expression of core and U1-specific spliceosomal proteins in the brains of Tau transgenic flies (*elav>Tau*^*R406W*^). We focused on 1-day-old adults preceding the onset of significant neuronal loss (Wittmann et al., 2001). Based on western blots prepared from adult head homogenates, *elav>Tau*^*R406W*^ flies demonstrate an approximately 30-40% reduction in the levels of SmB, SmD2, and SNF proteins compared to *elav-GAL4/+* controls (Figure 3A-B). Moreover, we did not detect any significant changes in mRNA levels based on quantitative real-time polymerase chain reaction (qRT-PCR), consistent with a post-transcriptional mechanism for the observed reductions (Figure 3C). In addition, immunofluorescence staining of adult brains using the anti-Sm antibody (Y12), which recognizes both SmB and SmD3 in *Drosophila* (Brahms et al., 2000; Gonsalvez et al., 2006) or an anti-SNF antibody confirmed markedly reduced U1 snRNP levels and depletion from neuronal nuclei (Figure 3D).

**Figure 3.**
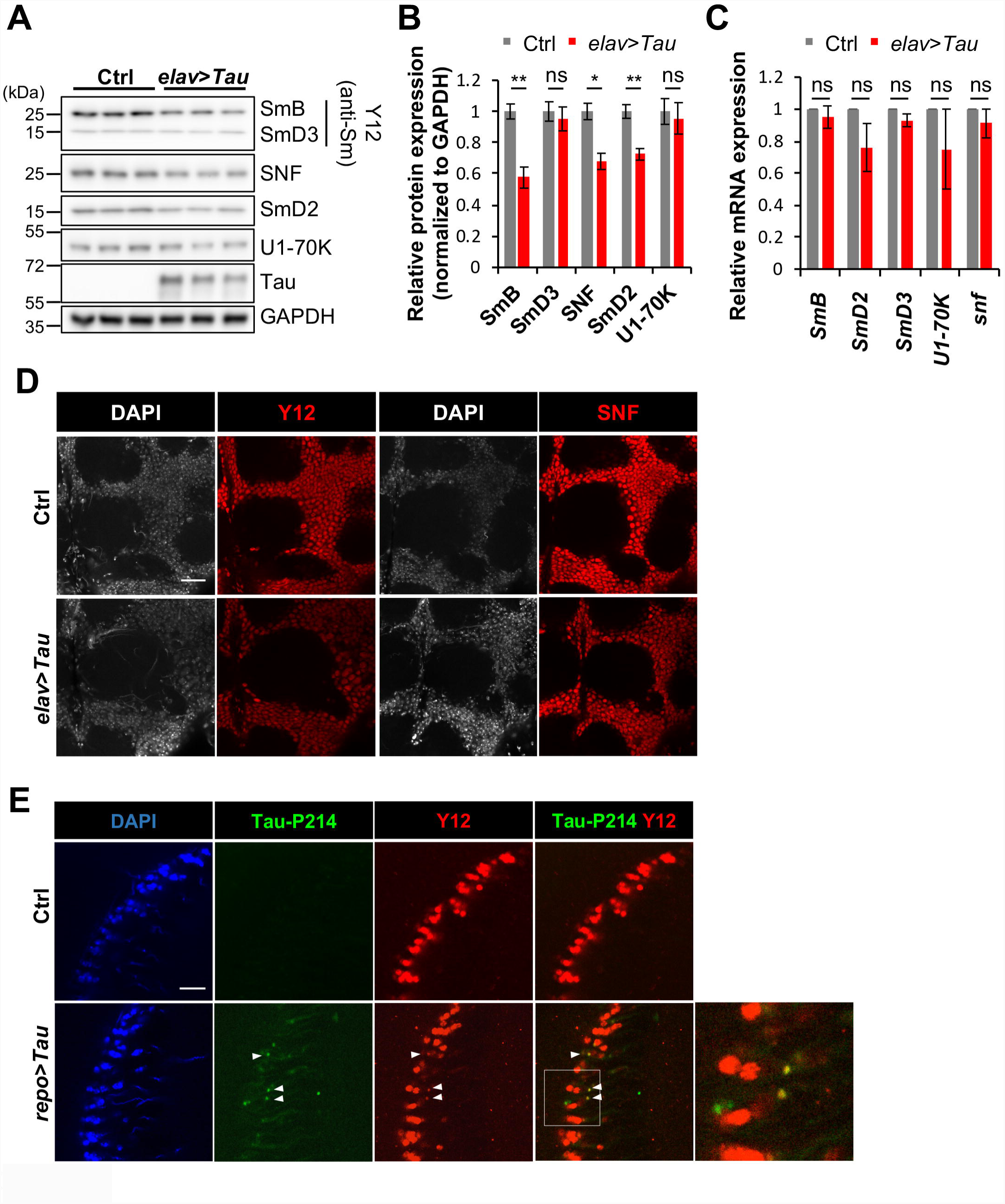
Disruption of the core spliceosome following Tau expression in *Drosophila*. (**A**) Pan-neuronal expression of human Tau causes reduced levels of multiple spliceosomal proteins. Adult fly head homogenates were prepared from 1-day-old *elav>Tau* (*elav*-*GAL4*/+; +/+; *UAS*-*Tau*^*R406W*^/+) or control animals (*elav*-*GAL4*/+), and western blots were probed for SmB, SmD3, SNF (U1A), SmD2, U1-70K, Tau and GAPDH (loading control). The anti-Sm antibody (Y12) recognizes both SmB and SmD3. (**B**) Quantification of spliceosomal protein levels from (**A**), normalized to GAPDH. SmB, SNF, and SmD2 exhibited significant reductions following Tau expression.(**C**) Pan-neuronal Tau expression does not significantly affect *SmB, SmD3, SmD2, U1-70K*, or *snf* mRNA levels based on quantitative real time-polymerase chain reaction (qRT-PCR). mRNA was prepared from 1-day-old adult heads, using the same genotypes as in **A**-**B**. (**D**) Whole-mount stains of adult fly brains reveal consistent depletion of SmB/D3 and SNF nuclear protein, using either the Y12 or anti-SNF antibodies, respectively (Red). Nuclei are stained with DAPI (Gray). Representative images of 1-day-old animals are shown, using the same genotypes as above. Scale bar: 20 μm. (**E**) Glial expression of human Tau, using the *repo-GAL4* driver, leads to aberrant cytoplasmic foci (arrowheads) of SmB/D3 (Y12, Red) that colocalize with tangle-like aggregates of phospho-Tau (anti-Tau-P214, Green) in 10-day-old adult brains. Nuclei are stained with DAPI (Blue). Representative images from control (*repo-GAL4, tub-GAL80*^*TS*^/+) and *repo*>*Tau* (*repo-GAL4, UAS*-*Tau*^*WT*^, *tub-GAL80*^*TS*^/+) are shown. Scale bar: 10 μm. See also Figure S3A for staining of 1-day-old animals and biochemical studies. For statistical analyses (**B** & **C**), unpaired t-tests were performed, based on 3 replicate experiments. *, p < 0.05; **, p < 0.01; ns, not significant. All error bars denote mean ± SEM.

In human AD postmortem brain tissue, multiple spliceosomal proteins can be found mislocalized to the cytoplasm, co-aggregating with Tau in neurofibrillary tangles (Bai et al., 2013; Hales et al., 2014). In *Drosophila* neurons, although Tau is misfolded and hyper-phosphorylated as in human AD, it remains predominantly soluble and oligomeric (Ali et al., 2012; Cowan et al., 2010; Mudher et al., 2004; Wittmann et al., 2001). In other human tauopathies, such as corticobasal degeneration, fibrillar Tau inclusions are observed in both neurons and glia, and prior work has established that Tau aggregates more readily when expressed in *Drosophila* glia, forming insoluble, tangle-like, cytoplasmic inclusions (Colodner and Feany, 2010). As a complementary approach, we therefore stained for Sm proteins in a *Drosophila* glial tauopathy model, which relies on the *repo-GAL4* glial driver. Indeed, aged *repo>Tau*^*WT*^ flies manifest numerous cytoplasmic aggregates co-staining for both phospho-Tau (anti-pSer214) and SmB/SmD3 (Y12) (Figure 3E). Consistent with this, on western blots, we can detect both Tau and increased SmB protein in insoluble fractions prepared from *repo>Tau*^*WT*^ heads (Figure S3B). Together, our results suggest that soluble Tau species lead to a loss of snRNP protein levels, whereas insoluble forms of Tau co-aggregate with spliceosomal proteins, leading to cytoplasmic sequestration.

### Spliceosome loss-of-function causes neurodegeneration

Our data suggest that pathological forms of Tau can trigger a reduction in core and U1-specific spliceosomal components in *Drosophila* neurons, and that further experimental reduction of these proteins enhances Tau-induced neurodegeneration. Therefore, we next examined whether disruption of the U1 snRNP is sufficient to cause neurodegeneration, independent of transgenic Tau. The spliceosome is essential for organismal development and maintenance of cellular functions. Available null or strong hypomorphic alleles for U1 snRNP components including *snf/U1A, U1-70K*, and *SmB* are embryonic lethal (Anne, 2010; Flickinger and Salz, 1994; Salz et al., 2004), hindering studies in the adult nervous system. *Drosophila SmB* is the single fly ortholog for both human *SNRPB* and *SNRPN*, which substitutes for SmB in neurons (McAllister et al., 1988; Saltzman et al., 2011). While attempting to generate a GFP-tagged allele of fly *SmB*, we serendipitously created a viable, hypomorphic allele. *SmB*^*MI07584*^ contains a *Minos*-mediated integration cassette (MiMIC) transposable element insertion within the first intron of *SmB* (Venken et al., 2011) (Figure 4A). Using recombinase mediated cassette exchange, the MiMIC element was replaced with a GFP coding exon flanked by splice acceptor/donor sequences (Nagarkar-Jaiswal et al., 2015a, 2015b). Hereafter, we refer to this allele as *SmB*^*MG*^ for <u>M</u>iMIC-<u>G</u>FP. The *SmB*^*MG*^ allele, encoding a full-length, N-terminal GFP-tagged SmB protein, is homozygous viable. In adult brains, the SmB^MG^ fusion protein is expressed at comparable levels to SmB in wild-type controls (Figure 4B), and is localized to the nucleus, as expected (Figure 4C). Surprisingly however, *SmB*^*MG*^ fails to complement several available *SmB* loss-of-function alleles, including *SmB*^*MI07584*^, *SmB*^*BG02775*^, *SmB*^*SH0509*^ or a deficiency strain covering the locus (*Df(2L)BSC453*, Figure 4A), causing embryonic lethality in compound-heterozygotes (Table S5). Viability is fully rescued by a 90-kb bacterial artificial chromosome (BAC) transgenic construct including the *SmB* genomic locus (*SmB*^*GR*^, Figure 4A). These data suggest that *SmB*^*MG*^ is a viable, hypomorphic allele encoding an SmB protein with reduced function, yet sufficient for embryonic development and adult viability.

**Figure 4.**
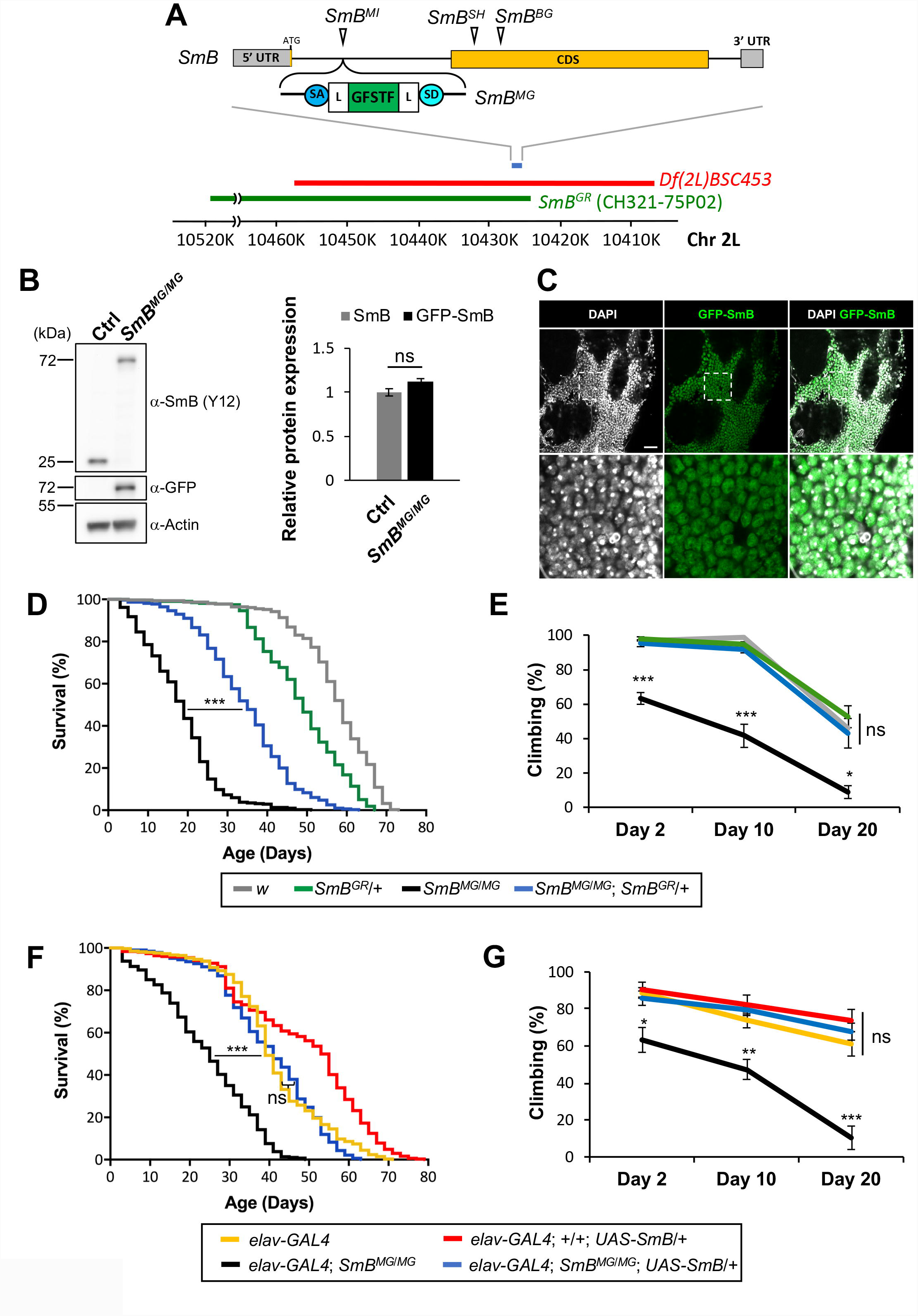
Loss of function in *SmB*, encoding a core spliceosome factor, causes reduced survival and progressive locomotor impairment. (**A**) The *Drosophila SmB* locus on chromosome 2L, including the single coding exon (CDS, Yellow) and untranslated regions (UTR, Gray) is depicted. Several transposable element insertion alleles used in this study are also noted including *SmB*^*MI*^ (*Mi{MIC}SmB*^*MI07584*^), *SmB*^*SH*^ (*P{lacW}SmB*^*SH0509*^), and *SmB*^*BG*^ (*P{GT1}SmB*^*BG02775*^). *SmB*^*MI*^ consists of a *Minos*-mediated integration cassette (MiMIC) inserted within the first intron. To generate the *SmB*^*MG*^ allele, recombination mediated cassette exchange was performed, introducing a coding exon for the green fluorescence protein (GFSTF: EGFP-FlAsH-StrepII-TEVcs-3xFlag), flanked by flexible linker (L) domains, and both splice acceptor (SA) and splice donor (SD) sequences. A chromosomal deficiency strain (Red), *Df(2L)BSC453*, deletes ∼51 kb including the entire *SmB* locus. The transgenic genomic rescue strain, *SmB*^*GR*^, carries a ∼90 kb Bacterial Artificial Chromosome (BAC, CH321-75P02, Green), including the *SmB* locus. See also Table S5. (**B**) *SmB*^*MG*^ homozygotes (*SmB*^*MG*/*MG*^) demonstrate expression of the GFP:SmB fusion protein (GFP-SmB) in *SmB*^*MG*^ flies at levels comparable with controls (*w*). Western blots from adult fly head homogenates were probed with anti-Sm (Y12), anti-GFP, or anti-Actin (loading control). Relative expression of SmB, normalized to Actin, was quantified based on analysis of 3 replicate experiments. (**C**) GFP-SmB (Green) is localized to the nucleus (DAPI, Gray) in brains from *SmB*^*MG*^ homozygous adults. Scale bar: 20 μm. Boxed region of interest in top row is shown magnified in bottom row. (**D**) *SmB*^*MG*/*MG*^ adults (Black) exhibit reduced survival compared with controls (Gray: *w*), and this phenotype is partially rescued by the *SmB* BAC transgenic (Blue: *SmB*^*MG*/*MG*^; *SmB*^*GR*^/+). Survival curve for *SmB*^*GR*^/+ (Green) is also shown. At least 313 adult flies were examined for each genotype. See also Figure S4A. (**E**) *SmB*^*MG*/*MG*^ adults also demonstrate progressive locomotor impairment, based on the startle-induced negative geotaxis response, and this phenotype is fully rescued by the *SmB* BAC transgenic. Genotypes are the same as in **D**. At least 5 replicate experiments were performed, each consisting of 7-18 flies, evaluating the proportion of flies climbing at least 5 cm in 5 sec. See also Figure S4B. (**F** and **G**) Both the *SmB*^*MG*/*MG*^ survival (**F**) and locomotor (**G**) phenotypes were rescued by tissue-specific expression of wild-type *SmB* in the nervous system, using the *elav*-*GAL4* pan-neuronal driver. The following genotypes were evaluated: *elav-GAL4*/*Y* (Yellow); *elav-GAL4*/*Y*; +/+; *UAS-SmB*/+ (Red); *elav- GAL4*/*Y*; *SmB*^*MG*/*MG*^ (Black); and *elav-GAL4/Y*; *SmB*^*MG*/*MG*^; *UAS-SmB*/+ (Blue). For survival analysis (**F**), at least 288 adult flies were examined per genotype. Survival of *elav-GAL4/Y*; *SmB*^*MG*/*MG*^; *UAS-SmB*/+ was not significantly different from *elav-GAL4*/*Y*; *SmB*^*MG*/*MG*^ controls, consistent with complete rescue. For locomotor assay (**G**), at least 4 replicate experiments were performed, each consisting of 5-15 adult flies. For statistical analyses, the unpaired t-test (**B**), Kruskal-Wallis test (**D** and **F**) followed by Dunn’s test for post hoc comparisons, and one-way ANOVA (**E** and **G**) with Tukey’s test for post hoc comparisons were performed. *, p < 0.05; **, p < 0.01; ***, p < 0.001; ns, not significant. All error bars denote mean ± SEM.

We first examined survival and locomotor behavior in *SmB*^*MG*^ adults, since these phenotypes have previously been associated with neurodegeneration, including in Tau transgenic flies (Lessing and Bonini, 2009; Mudher et al., 2004; Wittmann et al., 2001). Indeed, *SmB* loss-of-function caused reduced survival (Figure 4D and S4A) and age-dependent, progressive locomotor impairment, based on the startle-induced, negative geotaxis assay (climbing) (Figure 4E and S4B). These *SmB*^*MG*^ phenotypes were rescued by the *SmB* genomic construct (*SmB*^*MG*^; *SmB*^*GR*^/+), establishing specificity. In addition, the *SmB*^*MG*^ survival and locomotor phenotypes were rescued by pan-neuronal expression of a wild-type *SmB* cDNA (*elav>SmB*), suggesting that these phenotypes arise from reduced *SmB* function in neurons (Figure 4F-G). In sum, our data indicate that SmB is required for maintenance of nervous system function and survival with aging.

Next, we investigated for more direct evidence of neurodegeneration in *SmB*^*MG*^ adult brains. Hematoxylin and eosin stained paraffin sections revealed overall preserved adult brain morphology without overt evidence of neuropil vacuolar degenerative changes. We therefore examined specific, vulnerable cell populations previously reported in Tau transgenic (Bardai et al., 2018) and other *Drosophila* models of neurodegenerative disorders (Khurana et al., 2010; Merlo et al., 2014), including Kenyon cell and cholinergic neurons. Kenyon cells are interneurons in the *Drosophila* mushroom body, which is a hub for invertebrate learning and memory (Owald and Waddell, 2015). We discovered that Kenyon cell numbers declined with aging in *SmB*^*MG*^ flies (Figure 5A-C). Cholinergic neurons are particularly susceptible to loss in both human AD and in *Drosophila* Tau transgenic models (Coyle et al., 1983; Wittmann et al., 2001). We found that *SmB*^*MG*^ similarly causes progressive cholinergic neuronal loss in the fly lamina, based on quantification using a choline-acetyl transferase (*ChAT*)>*Beta-galactosidase* reporter (Figure 5D-F). Finally, similar to Tau transgenic flies (Dias-Santagata et al., 2007), we documented neuronal apoptosis using the TUNEL assay in *SmB*^*MG*^ flies (Figure S4C-E). Importantly, introduction of the *SmB* genomic construct rescued all of the observed *SmB*^*MG*^ neurodegenerative phenotypes. We also confirmed that RNAi-mediated knockdown of *SmB* causes consistent neurodegenerative phenotypes (Figure S5). Overall, our data suggest that loss-of-function of *SmB*, an essential, core spliceosome component is sufficient to cause age-dependent neurodegeneration and progressive nervous system dysfunction.

**Figure 5.**
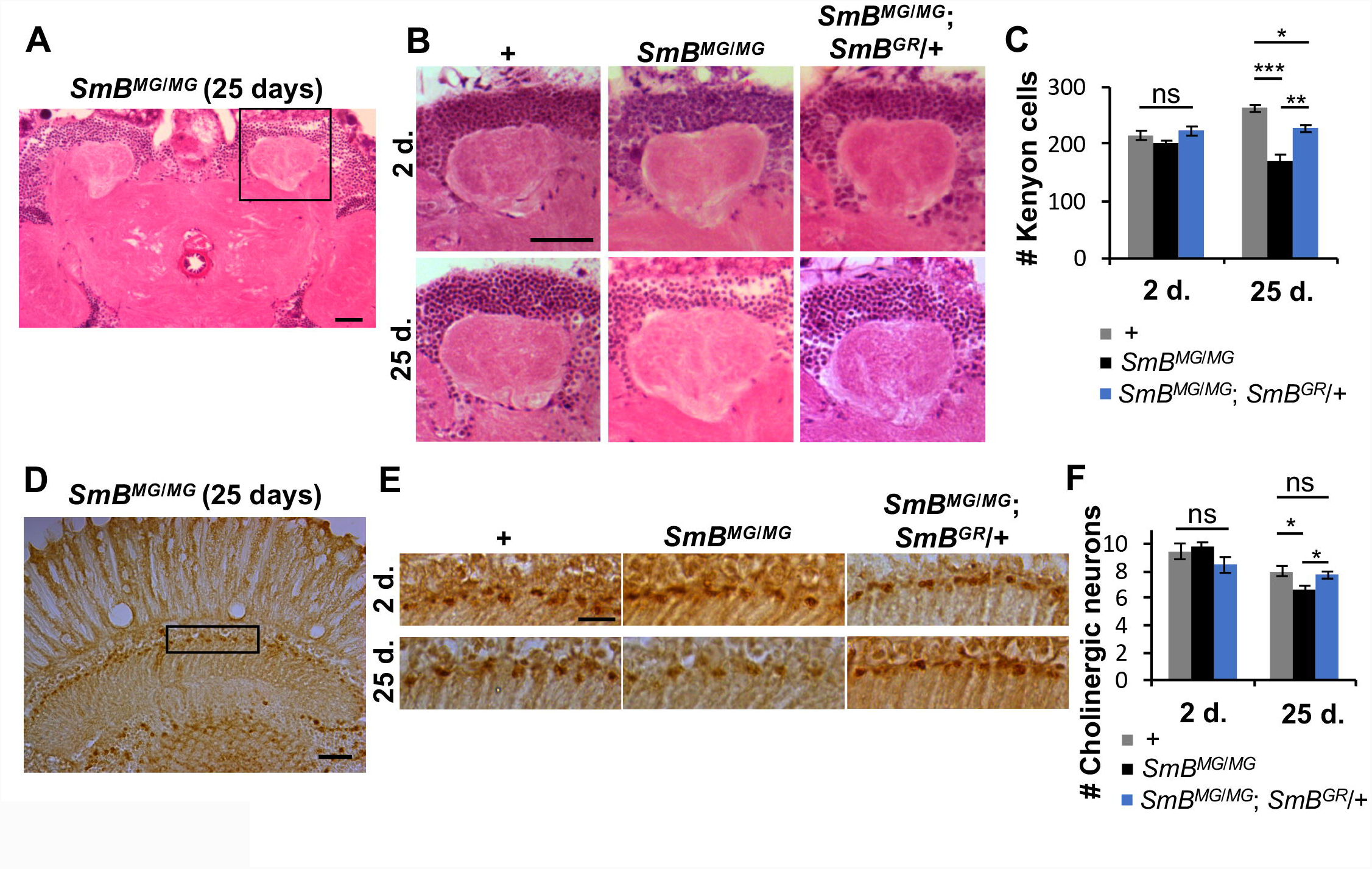
Loss of function in *SmB* causes neurodegeneration. (**A-C**) Loss of function in *SmB*, encoding a core spliceosome protein, causes age-dependent neuronal loss in adult brains. (**A**) Hematoxylin and eosin stained frontal brain section at the level of mushroom body calyces, highlighting the region of interest including the Kenyon cells. Scale bar: 25 μm. (**B**) Compared to controls (*w* and *SmB*^*MG*/*MG*^; *SmB*^*GR*^/+), *SmB*^*MG*/*MG*^ homozygous adults show progressive loss in Keynon cell numbers between 2- and 25-days (d.) of age. Scale bar: 25 μm. (**C**) Quantitation of experiment shown in **B**, based on Kenyon cell counts from at least 5 brains per genotype and timepoint. (**D-F**) *SmB* loss-of-function also causes progressive loss of cholinergic neurons in the lamina. (**D**) Frontal section at the level of lamina highlighting region of interest. Cholinergic neurons are labeled by DAB staining using the *ChAT*-*GAL4* reporter line driving expression of β-galactosidase (β-gal). Scale bar: 20 μm. (**E**) Compared to controls (*ChAT-GAL4*/*UAS-lacZ* and *ChAT-GAL4, SmB*^*MG*^/*UAS-lacZ, SmB*^*MG*^; *SmB*^*GR*^/+), *SmB* homozygotes (*ChAT-GAL4, SmB*^*MG*^/*UAS-lacZ, SmB*^*MG*^) exhibit loss of cholinergic neurons between 2- and 25-days (d.) of age. Scale bar: 10 μm. (**F**) Quantification of experiment shown in **E**, based on β-gal-positive neuronal counts from at least 9 animals for each genotype and timepoint. For statistical analyses (**C** and **F)**, one-way ANOVA followed by Tukey’s test for post hoc comparisons were performed. *, p < 0.05; **, p < 0.01; ***, p < 0.001; ns, not significant. All error bars denote mean ± SEM. See also Figure S4C-E and S5.

### Tau and spliceosome disruption cause similar splicing signatures in *Drosophila* brains

To determine if Tau is sufficient to cause splicing errors, possibly via interactions with the spliceosome, we performed RNA-sequencing (RNA-seq) on heads from Tau transgenics (*elav*>*Tau*^*WT*^ and *elav*>*Tau*^*R406W*^) or controls (*elav*-*GAL4*/*+*). Analyses were performed at 1-, 10-, or 20-days, to assess for potential changes relative to onset and progression of neurodegeneration. To examine the consequences of direct spliceosomal loss-of-function, RNA-seq was also performed on 10-day-old *SmB*^*MG*^ flies. We initially evaluated alternative splicing changes using the replicate Multivariate Analysis of Transcript Splicing (rMATS) tool (Shen et al., 2014). Indeed, we discovered up to 1,559 significant (false discovery rate [FDR] < 0.05), differential splicing events triggered by Tau expression in the adult fly brain (Figure S6A, Table S6, and Supplemental File 2). Tau-associated splicing changes were present in 1-day-old animals, preceding onset of neurodegenerative phenotypes, and were enhanced in flies expressing *Tau*^*R406W*^, a mutant form causing familial frontotemporal dementia. Genetic disruption of the core spliceosomal factor, *SmB*, was associated with even stronger transcriptome perturbations, causing 16,424 differential splicing events in 10-day old animals (Figure S6A, Table S6, and Supplemental File 2).

These results are consistent with our hypothesis that Tau-spliceosome interactions may significantly alter the transcriptional landscape. However, to differentiate aberrant splicing errors from alternative splicing we next applied two complementary analytic tools, including (1) a newly developed Differential Expression of Introns (DEIn) algorithm and (2) the previously-validated tool, CrypSplice (Tan et al., 2016). DEIn uses a stringent definition of intron retention as transcribed sequences mapping to gene loci, but completely absent in the *Drosophila* transcriptome reference (Figure S6B), consistent with aberrant suppression of otherwise, constitutively-spliced introns. By contrast, most intron retention events recognized by rMATS are documented alternative exons in annotated transcript isoforms. Similarly, CrypSplice facilitates sensitive detection of recurrent, cryptic splice junctions that are not present in annotated transcriptome references, causing shifts in either splice donor (5-prime) or acceptor (3-prime) positions, or both, as well as new combinations of splice donors and acceptors (Figure 6A). For this work, the CrypSplice software was further enhanced to improve annotation of cryptic splicing errors to facilitate interpretation of both causal mechanisms and potential consequences (see below and Methods). Indeed, our analyses identify a substantial number of splicing errors in Tau transgenic flies, including up to 437 cryptic splice junctions and 1,138 intron retention events (Figure 6B, Table S7-S10). As with the differential splicing analysis (above), splicing errors were detectable in 1-day-old animals prior to the manifestation of neurodegeneration and were more frequent in *Tau*^*R406W*^ than *Tau*^*WT*^ at all timepoints. Cryptic splicing errors also increased with aging at successive timepoints (Figure 6B). For each class of splicing error, we successfully validated several selected DEIn and CrypSplice predictions by RT-PCR (Figure 6C and Figure S6C).

**Figure 6.**
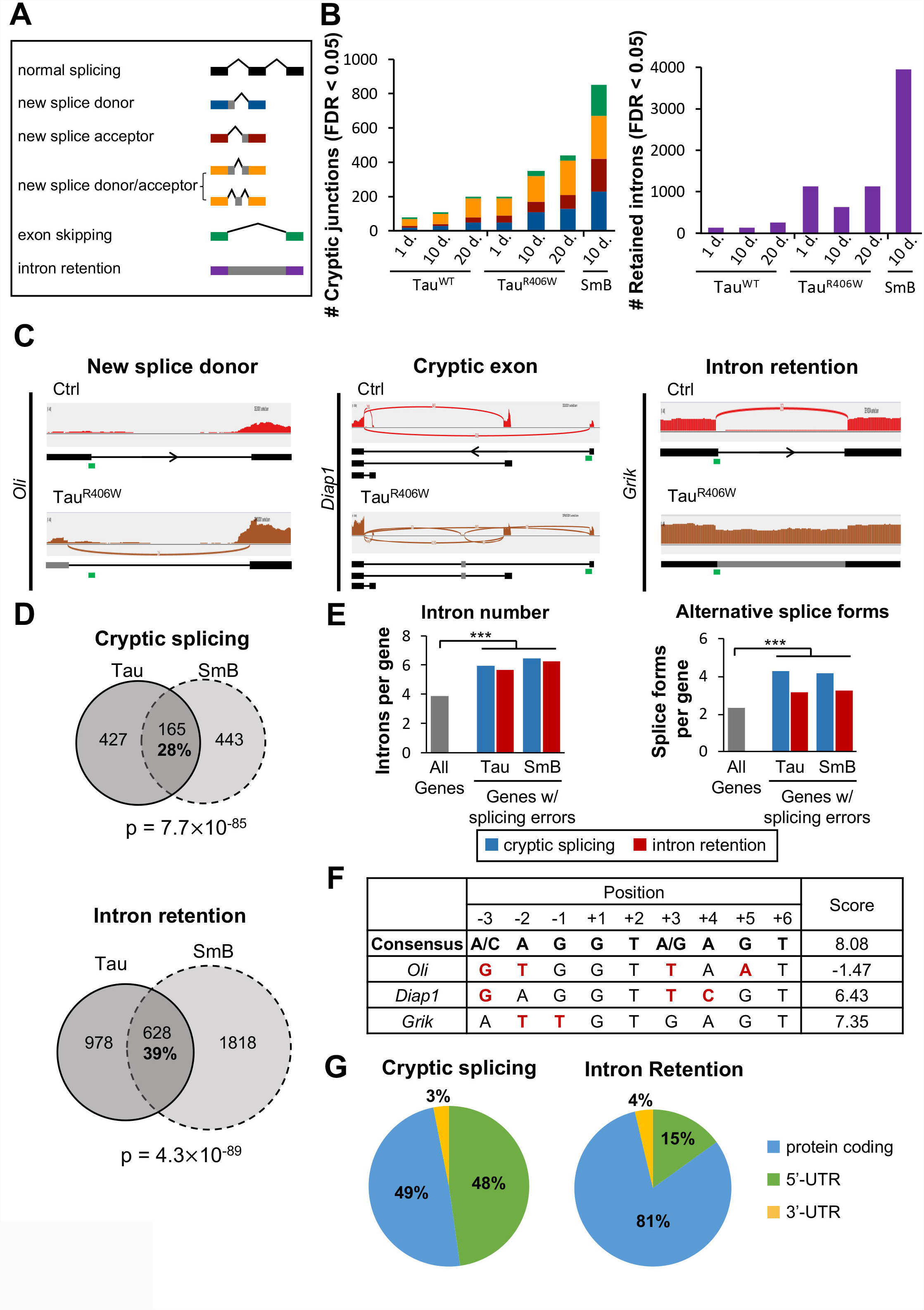
Tau expression and *SmB* loss cause similar profiles of splicing errors in *Drosophila* brains. (**A**) Types of splicing errors identified by CrypSplice and DEIn. (**B**) Pan-neuronal expression of Tau or *SmB* loss-of-function cause significant, differential expression of cryptic splicing errors (Left) and intron retention (Right). Cryptic splice junctions and retained introns were detected using the CrypSplice or DEIn tools, respectively, analyzing RNA-seq from *elav>Tau*^*WT*^ (*elav*-*GAL4*/+; *UAS*-*Tau*^*WT*^/+), *elav>Tau*^*R406W*^ (*elav*-*GAL4*/+; +/+; *UAS*-*Tau*^*R406W*^/+) or control animals (*elav*-*GAL4*/+), aged to 1-, 10-, or 20-days (d.). RNA-seq from *SmB*^*MG*/*MG*^ adults was compared to that from *y w* controls, both at 10 d. For statistical analyses of recurrent splicing errors, the beta binomial test was performed, followed by the Benjamini-Hochberg procedure for multiple test adjustment, with significance threshold set to FDR < 0.05. RNA-seq was performed on triplicate experimental samples (n = 3) for all genotypes, except for the control comparison for *elav>Tau*^*R406W*^, for which same batch, duplicate *elav*-*GAL4*/+ samples were used (n = 2) (see Methods). See also Figure S6B and Tables S7-10. (**C**) Plots highlighting representative Tau-induced splicing errors, including examples of a new splice donor (*Oli*, Left), a cryptic exon (*Diap1*, Middle), and intron retention (*Grik*, Right), based on analyses of 20 d. *elav>Tau*^*R406W*^ versus controls. Histograms denote the depth of RNA-seq read coverage, and curves denote split reads spanning splice junctions. The predicted resulting transcripts are indicated below, highlighting normal exonic structure (Black) from reference annotations versus aberrant exonic structure (Gray) resulting from non-annotated cryptic splice junction usage. Arrows denote the transcript orientation, and green bars indicate the annotated 5’ splice donor sites which were analyzed in **F**. See Figure S6C for experimental confirmation by RT-PCR. (**D**) The genes harboring splicing errors following pan-neuronal expression of Tau strongly overlap with that seen following *SmB* loss-of-function. We considered the union of all genes with significant, differential expression of cryptic splice junctions or intron retention, detected at 1, 10, or 20 d. in *elav*>*Tau*^*R406W*^ animals or from complementary analyses of 10 d. *SmB*^*MG*/*MG*^ flies; these gene sets were also used for analyses in **E** and **G**. Gene counts affected by cryptic splicing errors (Top) or intron retention (Bottom) are shown, along with the percent of Tau-associated, differentially-spliced genes that overlap. For statistical analysis, the hypergeometric overlap test was performed. (**E**) Splicing errors occur more commonly in genes with greater numbers of introns (Left) or annotated alternatively spliced transcripts (Right). Average gene/transcript feature (introns or spliceforms, respectively) shown for all *Drosophila* genes versus gene sets affected by cryptic splicing (Blue) or intron retention (Red) in *elav*>*Tau*^*R406W*^ or *SmB*^*MG*/*MG*^ animals. For statistical analysis, an empirical p-value was computed for each comparison based on random sampling (1000 iterations) from all *Drosophila* genes, using a sample size equal to the experimentally-defined gene sets with splicing errors in *Tau* or *SmB* animals. ***, p < 0.001. (**F**) Splicing errors occur at exon/intron junctions with splice donor sequences that diverge from the consensus motif for U1 spliceosome binding. The *Drosophila* consensus splice donor (5’ splice site) sequence (boldface) is shown, along with splice donor sequences corresponding to splicing errors in representative genes (Green bars in panel **C**). Nucleotides that diverge from the consensus are shown (Red), along with the corresponding splice site binding strength score calculated using MaxEntScan. See also Figure S6D. (**G**) Splicing errors are likely to disrupt protein expression. Pie charts show the frequency of Tau-associated cryptic splicing (Left) or intron retention errors (Right) affecting transcript coding sequences (Blue) or untranslated regions (UTR) (Green: 5’-UTR; Yellow: 3’-UTR). See also Tables S11 & S12.

To better understand the underlying mechanism(s), we next examined the profile of splicing errors detected in *Tau*^*R406W*^ transgenic flies, pooling results from all 3 time points and focusing on 592 genes harboring 985 cryptic splicing errors and 1,606 genes with 2,767 retained introns (Tables S11 & S12). Our analyses of *SmB* loss-of-function animals revealed comparable numbers of recurrent splicing errors (608 or 2,446 genes affected by cryptic splicing or intron retention, respectively) (Figure 6B, Table S7-S10). In fact, we found a 28% overlap in the specific genes affected by cryptic splicing (39% overlap for intron retention), and these results were significantly greater than that expected due to chance (p_cryptic_ = 7.7×10^−85^ or p_intron_ = 4.3×10^−89^, Figure 6D). Compared to all *Drosophila* genes, those vulnerable to splicing errors had significantly increased numbers of introns and more alternative splice forms (p<0.001, Figure 6E). Tau-induced cryptic splicing errors affected either the splice donor and/or acceptor sites (Figure 6B). Based on our experimental data suggesting that several spliceosomal proteins may be reduced and therefore dose-limiting in Tau transgenic flies (Figure 3), we hypothesized that splicing errors might occur preferentially at annotated splice donor/acceptor sites that diverge from the U1 or U2 consensus binding sites, respectively. For this analysis, we limited our consideration to those cryptic junctions for which the originally annotated, apparently skipped, donor or acceptor sequences could be inferred unambiguously (see Methods). Indeed, we found that the average estimated binding strength for these “error-prone” splice donor and acceptor sites was significantly lower (p<0.001) than for all annotated sites in the *Drosophila* genome (Figure 6F and Figure S6D). Consistent results were also obtained for splice junctions flanking Tau-triggered intron retention events. Overall, our data suggest that Tau-induced splicing errors have a similar profile to *SmB* loss-of-function, consistent with a shared mechanism due to spliceosomal disruption.

Lastly, to understand the potential functional consequences, we further classified splicing errors based on their potential to disrupt protein expression. Indeed, 49% of cryptic splicing errors and 81% intron retention events detected in Tau transgenic flies overlap with coding exons, with concomitant potential for open reading frame disruption (Figure 6G). Most of the remaining splicing errors are predicted to affect 5’ UTR sequences and may therefore also impact mRNA stability and/or translation. Based on enrichment analysis for gene ontology (GO) terms, Tau-induced cryptic splicing errors affected genes with predominant roles in protein phosphorylation (p = 4.8×10^−6^), synaptic vesicle exocytosis (p = 3.4×10^−4^), and neurotransmitter transport (p = 3.4×10^−4^), whereas Tau-induced intron retention affected genes implicated in the innate immune response (p = 1.0×10^−10^) and oxidation-reduction (p = 1.8×10^−5^) (Table S13). Overall, our data suggest that Tau-induced splicing errors likely have broad impact on the *Drosophila* brain transcriptome—with potential consequences for CNS function/maintenance—and cause a profile similar to genetic disruption of *SmB*, encoding a core spliceosomal factor.

### Tau pathology is associated with cryptic splicing errors in human brains

Recent studies of human postmortem brain have identified mRNA splicing changes, including intron retention, in the setting of AD pathology (Bai et al., 2013; Raj et al., 2018). While evidence of cryptic splicing has been identified in ALS (Humphrey et al., 2017; Ling et al., 2015; Tan et al., 2016), to our knowledge, this class of splicing error has not previously been examined in the context of AD. To determine if Tau neuropathology in AD is associated with cryptic splicing errors, we leveraged data from the Religious Orders Study and Rush Memory and Aging Project (ROSMAP) (Bennett et al., 2018; Mostafavi et al., 2018). Our analyses included 620 deceased subjects with comprehensive clinical and pathologic characterization (Table S14) and RNA-seq profiling of the dorsolateral prefrontal cortex (De Jager et al., 2018). As in our *Drosophila* analyses, we first implemented CrypSplice, examining for cryptic splice errors among 241 control versus 379 AD cases, based on AD consensus pathologic diagnostic criteria (see Methods). We identified few changes (n = 14) meeting our significance threshold (FDR < 0.05, Table S15). In a complementary analysis focusing on 100 cases and 100 controls with high or low Tau pathologic burden, respectively, we identified a modestly increased number of cryptic splicing errors (n = 56) between the 2 groups (Table S16). Compared to our studies in *Drosophila* models, we reasoned that inter-individual heterogeneity may prevent detection of cryptic splicing events that recur among such a large number of samples. Moreover, if Tau-induced cryptic splicing errors occur stochastically and at low-frequency, they may be distributed widely throughout the transcriptome, such that few recurrent errors might be detected among hundreds of samples. We therefore considered an alternative approach, deriving a person-specific “Cryptic Load” score, based on the average strength of cryptic junctions (See Methods). The Cryptic Load algorithm has been incorporated into our extended version of CrypSplice, enhancing this software tool for analysis of large scale RNAseq datasets derived from human population data, such as ROSMAP. Regression was first performed using consensus AD pathologic diagnosis (n cases/controls = 379/241) as an outcome, revealing a modest but non-significant increase in cryptic load in association with AD pathology (β = 1.7, p = 0.09, Table 1), after adjustment for age at death, postmortem interval (PMI), and sample batch. To improve our statistical power to detect Tau-associated cryptic splicing errors, we next considered an alternative strategy in which a subset of subjects were dichotomized into high (n = 136) versus low (n = 105) neurofibrillary tangles, based on Braak staging consensus criteria (Braak and Braak, 1991). We observed a significant association between Tau and cryptic load (β = 2.6, p = 0.01). Lastly, we employed a quantitative measure of neurofibrillary tangle burden as the outcome, and performed a sensitivity analysis, in which subjects were selected from each tail of the distribution (high versus low tangle burden) and examined for differences in cryptic load using regression. Consistent with our hypothesis, we found an increased estimate effect size and significant associations between Tau pathologic burden and cryptic load for more extreme comparisons among nested case/control comparisons considering total sample sizes from 600 down to 100 brains (Table 1 and Figure S6E). In sum, our data support a conserved relationship between Tau pathologic burden and cryptic splicing errors in human brains.

**Table 1.** Analyses of cryptic load in human postmortem brains.

| **Predictor Variable** | **Sample size** | **Coefficient** | **p-value** |
| --- | --- | --- | --- |
| AD diagnosis<sup>1</sup> | 620 (241 vs. 379) | 1.7 | 0.09 |
| Braak (high/low)<sup>2</sup> | 241 (105 vs. 136) | 2.6 | 0.01 |
| Tangle burden<sup>3</sup> | 600 (300 vs. 300) | 1.6 | 0.1 |
|  | 400 (200 vs. 200) | 1.8 | 0.07 |
|  | 300 (150 vs. 150) | 2.2 | 0.03 |
|  | 200 (100 vs. 100) | 2.3 | 0.02 |
|  | 100 (50 vs. 50) | 3.0 | 0.004 |

<sup>1</sup>Pathologic AD based on NIA-Reagan
<sup>2</sup>High/Low Braak tangle score (Braak 0-2 vs. 5-6)
<sup>3</sup>Neurofibrillary tangle burden, nested case/control analysis considering the extremes of the trait distribution for all samples.

## DISCUSSION

Integrating data from human brain autopsy cohorts and *Drosophila* models, we discover a novel mechanism of Tau-mediated neurodegeneration in AD converging on mRNA splicing. First, we show that numerous spliceosome components are physically associated with Tau in human brains with AD pathology, and in *Drosophila*, genetic manipulation of these factors enhances Tau neurotoxicity. Second, we find that transgenic expression of human Tau causes a reduction of multiple spliceosome components, and loss-of-function of the core spliceosome protein, *SmB* is sufficient to induce progressive neuronal dysfunction and loss independent of Tau. Lastly, we show that Tau causes splicing errors in *Drosophila* similar to genetic disruption of the spliceosome, and we confirm increased cryptic splicing load in association with Tau pathologic burden in human postmortem brain. Overall, our data support a model (Figure 7) whereby Tau-spliceosome interactions disrupt snRNP function, leading to splicing errors, loss of transcriptome fidelity, and ultimately neurodegeneration.

**Figure 7.**
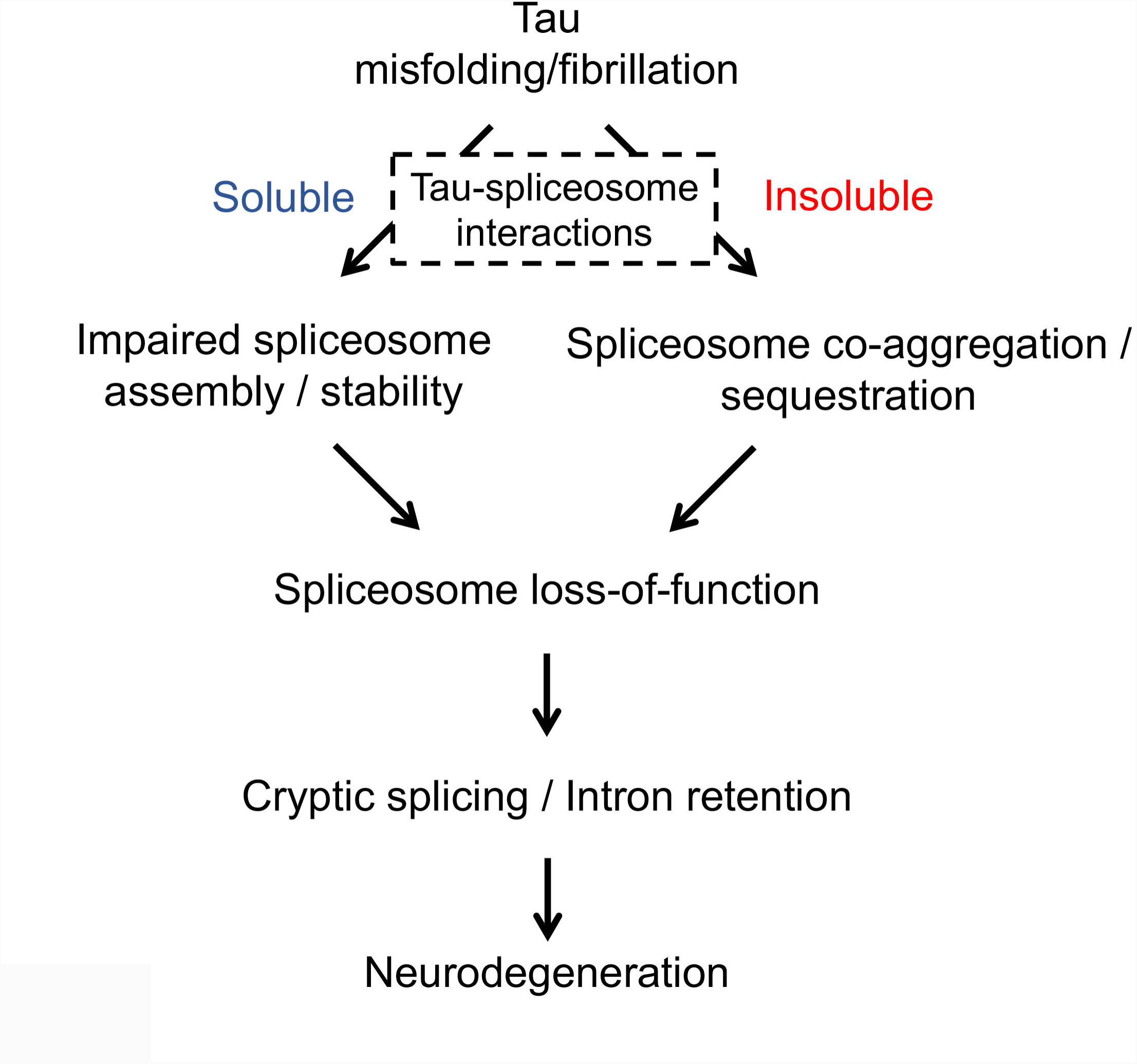
Hypothetical model for Tau-spliceosome interactions leading to neurodegeneration in AD. Based on evidence from human postmortem brain tissue and *Drosophila* transgenic models, we propose that multiple spliceosome components can associate with either soluble or insoluble toxic Tau species. In the case of insoluble Tau, spliceosome components that co-aggregate may be sequestered in the cytoplasm with neurofibrillary tangles. Alternatively, soluble Tau species can also interact with spliceosomal proteins, potentially disrupting snRNP assembly and/or stability, leading to reduced expression levels. In *Drosophila* models, knockdown of genes encoding core spliceosome components enhances Tau neurotoxicity. Furthermore, SmB loss-of-function causes a similar profile of cryptic splicing and intron retention as in Tau transgenic flies, and also causes age-dependent neurodegeneration. Therefore, Tau-spliceosome interactions likely contribute to Tau-associated CNS dysfunction and neuronal loss, possibly due to accumulated splicing errors and widespread transcriptome perturbation.

### The spliceosome, transcriptome integrity, and maintenance of the nervous system

In humans, dominantly-inherited mutations in *SNRPB* cause cerebro-costo-mandibular syndrome, a rare developmental disorder (Lynch et al., 2014). The *SNRPB* paralog, *SNRPN*, which is specifically expressed in the nervous system, is found at the imprinted, Prader-Willi syndrome locus, and altered expression has been implicated in the etiology of neurodevelopmental delay (Cassidy et al., 2012). Given the essential role of splicing for cellular function and embryogenesis, nearly all mutations previously recovered in core spliceosome proteins, including *SmB*, are embryonic lethal in *Drosophila* and other animal models. Our serendipitous discovery of a viable, hypomorphic allele of *SmB*, encoding a ubiquitously-expressed, core component of the spliceosome, facilitates study of the fundamental cellular splicing machinery in the maintenance and function of the adult nervous system. The brain appeared normally developed in adult *SmB* mutant flies; however, in aged animals we documented neuronal loss along with progressive locomotor impairment and decreased survival. Interestingly, these defects were rescued by neuronal-specific expression of wild-type *SmB*. Therefore, the *Drosophila* nervous system is especially vulnerable to reduction in spliceosome function, and susceptibility for resulting neurodegeneration increases with aging. Splicing is a major driver of transcriptome diversity, and in both *Drosophila* and mammals, alternative splicing is highest in the brain compared with all other tissues, consistent with an important role for the splicing machinery in neuronal diversity and brain health (Li et al., 2007; Raj and Blencowe, 2015). While additional studies will be required to confirm the precise mechanisms, it is likely that degradation of transcriptome fidelity in *SmB* mutant flies directly results in progressive neuronal dysfunction and death. RNA-seq profiles reveal that *SmB* loss-of-function causes massively dysregulated splicing, including thousands of differentially expressed splice forms and hundreds of splicing errors, including intron retention and cryptic junctions. These data are consistent with prior reports of genetic manipulation of spliceosome components in mouse models, in which aberrant pre-mRNA splicing is accompanied by neurodegeneration. For example, loss of either the *U2 snRNA* or *RBM17* induced cryptic splice junctions and intron retention, along with prominent cerebellar degeneration (Jia et al., 2012; Tan et al., 2016).

### Splicing errors and transcriptome fidelity in AD

Spliceosome disruption and altered pre-mRNA processing are also implicated in the pathogenesis of human diseases, including hematologic cancers (Hsu et al., 2015; Yoshida et al., 2011) and multiple neurologic disorders (Cooper et al., 2009). Mutation of *SMN*, encoding an essential cofactor for snRNP biogenesis, causes the recessive neurodegenerative disease, spinal muscular atrophy (Lefebvre et al., 1995; Lorson et al., 1999). More recently, genetic variants affecting numerous RNA-binding protein splicing factors have been identified in familial FTD-ALS (Ito et al., 2017), and most of these proteins, including TDP-43, FUS, TAF15, TIA1, and hnRNPA2B1, closely associate with the spliceosome complex (Förch et al., 2002; Freibaum et al., 2010; Leichter et al., 2011; Martinez et al., 2016; Sun et al., 2015).

In the *Drosophila* nervous system, Tau induced a similar profile of splicing errors as *SmB* loss-of-function, and genetic manipulation of numerous core snRNP components enhanced Tau toxicity. Many of the same genes were vulnerable to splicing errors in Tau transgenics and *SmB* mutants. Further, we found that the splice donor and acceptor sites most susceptible to errors had weaker binding sites for the U1 and U2 snRNPs, respectively. Mutant forms of Tau causing accelerated neurodegenerative phenotypes in both humans and flies also showed more profound transcriptome disruption, and cryptic splicing errors were more frequent in aged animals. Together, these data strongly suggest that spliceosome disruption and resulting splicing errors may mediate Tau-induced neurotoxicity in AD and other tauopathies. Compared to studies of FTD-ALS (Humphrey et al., 2017; Ling et al., 2015; Polymenidou et al., 2011; Tan et al., 2016), altered splicing has only recently been systematically investigated in AD. Raj et al. (2018) reported on global splicing changes associated with human cortical Tau pathologic burden, including hundreds of potential intron retention events. Consistent with our findings in *Drosophila* models, *MAPT* expression in human neuronal cultures was sufficient to trigger splicing errors. Using the same ROSMAP brain autopsy sample, we now extend these findings to demonstrate evidence of cryptic splicing errors in human tissue with AD pathology. To our knowledge, this class of error previously described in FTD-ALS (Humphrey et al., 2017; Ling et al., 2015; Tan et al., 2016) has not been reported in AD. The extended CrypSplice tool used for our study focuses exclusively on new, non-annotated junctions. Therefore, many of the Tau-associated changes that we highlight likely represent *bona fide* splicing errors, consistent with our hypothesis that Tau may disrupt the fundamental splicing machinery.

We argue that splicing errors are likely a cause rather than a consequence of Tau-induced neurotoxicity. First, splicing changes were detectable in young flies, preceding the onset of neurodegeneration (Wittmann et al., 2001). Second, our finding of dose-sensitive genetic interactions following reduction in spliceosome components suggests these factors are causal mediators of Tau toxicity. Third, direct manipulation of a core spliceosome component, *SmB*, causes neurodegeneration in *Drosophila*. Broadly, we propose 2 mechanistic models for how splicing errors and resulting transcriptome perturbation may promote neurodegeneration in AD. In the first, global degradation in transcriptome fidelity may overwhelm cellular RNA and/or protein quality control mechanisms (Garneau et al., 2007; Pilla et al., 2017). Accumulation of non-productive transcripts might be directly cytotoxic, and translation could also lead to misfolded, dysfunctional proteins and resulting proteostatic stress. Our recent study of AD postmortem brain tissue identified “cryptic peptides” corresponding to many mRNA splicing alterations, including those creating new exonic splice junctions (Johnson et al., 2018). In an alternate model, selected cellular pathways essential to neuronal health and/or survival may be particularly vulnerable to splicing errors. In fact, Tau-associated splicing errors were significantly enriched in genes implicated in synaptic function and immune response, which are each strongly implicated in AD pathogenesis (Heppner et al., 2015; Spires-Jones and Hyman, 2014), and similar processes were impacted by cryptic splicing errors in TDP-43 cellular and mouse models relevant to FTD-ALS (Polymenidou et al., 2011). These 2 models are not mutually exclusive, and it is possible that Tau-induced transcriptomic changes promote a global cellular stress response, while simultaneously targeting specific cellular pathways that hasten the demise of neurons.

### Tau-spliceosome interactions in AD

Additional work will be required to define the precise mechanism(s) by which Tau disrupts the spliceosome, but our findings and other published evidence provide important clues. One possibility is that insoluble Tau co-aggregates with and sequesters spliceosomal factors in the cytoplasm (Figure 7, right). In prior work, multiple core and specific components of the U1 snRNP were abnormally enriched with misfolded/aggregated Tau in insoluble protein fractions from AD postmortem brain, and these proteins closely associated with Tau in neurofibrillary tangles (Bai et al., 2013; Bishof et al., 2018; Hales et al., 2014; Johnson et al., 2018). However, abundant evidence suggests that Tau neurotoxicity may be mediated by predominantly oligomeric, soluble forms (Cowan and Mudher, 2013). Additionally, while Tau and spliceosomal factors co-aggregated in *Drosophila* glia, the sequestration model cannot easily account for all of our findings since Tau does not form substantial insoluble aggregates in fly neurons (Wittmann et al., 2001). We therefore hypothesize that soluble forms of Tau may also interact with and disrupt the assembly and/or stability of the spliceosome (Figure 7, left). Consistent with this, we discovered that Tau co-immunoprecipitates with spliceosomal proteins in soluble fractions from AD postmortem brain homogenates. Interestingly, similar data were recently reported in the mouse rTg4510 Tauopathy model, where numerous RNA-binding proteins co-localized with diffuse and oligomeric forms of Tau but were excluded from more mature, fibrillar tangles (Maziuk et al., 2018). Tau, along with many RNA-binding proteins, are intrinsically disordered proteins capable of liquid-liquid phase separation (Ambadipudi et al., 2017; Bishof et al., 2018; Kim et al., 2013; Lee et al., 2016; Molliex et al., 2015; Patel et al., 2015; Wegmann et al., 2018). Such interactions facilitate the assembly of “membrane-less organelles” with critical roles in RNA processing, such as cytoplasmic RNA stress granules and nuclear speckles, that regulate translation and splicing, respectively (Hyman et al., 2014; Lin et al., 2015; Nott et al., 2015). However, pathologic fibrillization of disease-associated proteins can perturb cellular dynamics of liquid-liquid phase separation with concomitant functional disruption of membrane-less organelles (Lee et al., 2016; Patel et al., 2015). For example, recent studies suggests that interactions between Tau and RNA-binding proteins, including ribosomal proteins and RNA stress granule components, can disrupt translation (Meier et al., 2016). Interestingly, hexanucleotide expansions in *C9ORF72*, the most common cause of FTD-ALS, trigger accumulation of cytotoxic dipeptide repeats that avidly bind U2 snRNP components and disrupt spliceosome assembly (Yin et al., 2017). In mammalian cells, failure to assemble the snRNP core, leads to rapid lysosomal degradation of Sm proteins (Prusty et al., 2017). Potentially consistent with this, we found that Tau induced a ∼30% reduction in multiple spliceosomal proteins in *Drosophila* brains. Tau has also recently been demonstrated to disrupt nuclear pore integrity and function (Eftekharzadeh et al., 2018); therefore, either altered assembly and/or nucleocytoplasmic transport of snRNPs might account for our findings. In sum, our data support defective spliceosomal biogenesis and function as a novel mediator of Tau-induced neurotoxicity, highlighting intriguing parallels with FTD-ALS, and further implicating downstream transcriptome perturbations as a contributor to neurodegeneration in AD.

## Supporting information

Supplemental File 2 (Differential Alternative Splicing dataset)

## ACKNOWLEDGEMENTS

We thank H.J. Bellen, J. Botas, M.B. Feany, A.G. Matera, H.K. Salz for generously providing antibodies and *Drosophila* stocks. We thank the Bloomington *Drosophila* Stock Center, the Vienna Drosophila RNAi Center, and FlyBase (Gramates et al., 2017). This study was supported in part by grants from the NIH (R01AG036836, R01AG050631, R01AG057339, U01AG046161, P30AG10161, R01AG15819, R01AG17917, U01AG46152, R01GM120033). J.M.S. was additionally supported by grants from Huffington Foundation, Jan and Dan Duncan Neurological Research Institute at Texas Children’s Hospital, and a Career Award for Medical Scientists from the Burroughs Wellcome Fund. Z.L. received additional support from the Cancer Prevention Research Institute of Texas RP170387, Houston Endowment and Belfer Neurodegenerative Disease Consortium. The Pathology and Histology Core at Baylor College of Medicine is supported by NIH grant P30CA125123. The work was additionally supported by U54HD083092 from the Eunice Kennedy Shriver National Institute of Child Health & Human Development. The results published here are in part based on data obtained from the AMP-AD Knowledge Portal accessed at doi:10.7303/syn2580853. We thank Drs. Thomas Cooper and Juan Botas for critical feedback and helpful discussions.

## AUTHOR CONTRIBUTIONS

Conceptualization, Y.-C.H., C.G., H.K.Y., M.A., N.T.S., Z.L., J.M.S.; Investigation, Y.-C.H., C.G., M.A., Y.L.; Methodology, H.K.Y., Z.L.; Formal Analysis, Y.-C.H., C.G., H.K.Y., R.A.-L., E.B.D.; Data Curation, H.K.Y., R.A.-L.; Writing—Original Draft, Y.-C.H., C.G., J.M.S.; Writing—Review & Editing, Y.-C.H., C.G., H.K.Y., M.A, R.A.-L., E.B.D, Y.L., J.A.L., A.I.L., D.A.B., P.L.D., N.T.S., J.M.S., Z.L.; Funding Acquisition, J.A.L., A.I.L., D.A.B., P.L.D., N.T.S., J.M.S., Z.L.; Resources, D.A.B, P.L.D.; Supervision, N.T.S, J.M.S., Z.L.

## METHODS

### Human subjects

The Religious Orders Study and Rush Memory and Aging Project (ROSMAP) participants were free of known dementia at enrollment, agreed to annual clinical evaluations, and signed an informed consent and Anatomic Gift Act donating their brains at death, approved by the Institutional Review Board at Rush University (Bennett et al., 2018). AD clinical diagnoses were made following National Institute of Neurological and Communicative Disorders and Stroke-Alzheimer’s Disease and Related Disorders Association recommendations (McKhann et al., 1984). Modified Bielschowsky silver stain was used to visualize neuritic plaques, diffuse plaques, and neurofibrillary tangles in tissue sections from the midfrontal, middle temporal, inferior parietal, and entorhinal cortices and the hippocampal CA1 sector (Bennett et al., 2006). AD neuropathologic diagnosis was made based on intermediate or high likelihood of AD by criteria from the National Institute on Aging and the Reagan Institute Working Group on Diagnostic Criteria for the Neuropathological Assessment of Alzheimer’s Disease (National Institute on Aging, 1997). Subjects were also classified based on Braak staging consensus criteria (Braak and Braak, 1991), including high (5-6) and low (0-2) neurofibrillary tangle pathology scores. As in prior work (Bennett et al., 2009), a quantitative composite score for neurofibrillary tangle pathologic burden was created by dividing the raw counts in each region by the population standard deviation of the region-specific counts and then averaging the scaled counts over the 5 brain regions to create a single standardized summary measure. Clinical and demographic features of the ROSMAP decedents included in this study can be found in Table S14.

## METHOD DETAILS

### Tau immunoprecipitation and human postmortem brain proteomics

Healthy control (n = 4) and AD patient (n = 4) frontal cortex brain samples (Table S1) were homogenized in NP-40 lysis buffer (25 mM Tris-HCl at pH 7.5, 150 mM NaCl, 1 mM EDTA, 1% NP-40, 5% Glycerol, 5 mM iodoacetamide) with protease and phosphatase inhibitors using a bullet blender according to manufacturer’s instructions followed by centrifugation at 10,000 x *g* for 15 min at 4 °C. Immunoprecipitation was done on 1 mg of protein from each sample as follows: brain lysates were first pre-cleared using Protein A-Sepharose conjugated beads (Invitrogen). Lysates were incubated with 3 μg of anti-Tau monoclonal antibody (TAU-5) overnight at 4 °C. Purified Mouse IgG2a K isotype (BD Pharmingen) was used as an isotype-matched negative control. 50 μL of Protein G DynaBeads (Invitrogen) beads were incubated with the lysate for 1 hr. Beads were then washed 3 times using wash buffer (50 mM Tris HCl at pH 8.0, 150 mM NaCl and 1% NP-40) and rinsed three times using phosphate buffered saline (PBS). A portion of the Dynabeads were subjected to western blot analysis (below) (Figure 1A).

The remaining beads were rinsed in PBS (3X) and re-suspended in 500 µL 50 mM NH_4_HCO_3_. Proteins bound to beads were reduced using 1 mM dithiothreitol (DTT) for 30 min and alkylated with 5 mM iodoacetamide (IAA) for 30 min in darkness. Proteins were digested with 1:100 (w/w) lys-C endopeptidase (Wako) at room temperature for 3 hr followed by further overnight digestion with 1:50 (w/w) trypsin (Promega) at RT. Tryptic peptides were acidified using 1% formic acid and 0.1% trifluoroacetic acid before desalting and purification using C18 StageTip. Peptides were eluted in 50% acetonitrile and were dried using a SpeedVac (Savant). Dried peptides were reconstituted in peptide loading buffer (0.1% formic acid, 0.03% trifluoroacetic acid, 1% acetonitrile). Peptide mixtures were separated by liquid chromatography on a self-packed C18 (1.9 um Dr. Maisch, Germany) fused silica column (25 cm × 75 μM internal diameter; New Objective, Woburn, MA) by a NanoAcquity UHPLC (Waters, Milford, FA) and monitored on a Q-Exactive Plus mass spectrometer (ThermoFisher Scientific, San Jose, CA). Elution was performed over a ∼120-minute gradient at a rate of 400 nL/min with buffer B ranging from 3% to 80% (buffer A: 0.1% formic acid and 5% DMSO in water, buffer B: 0.1 % formic and 5% DMSO in acetonitrile). The mass spectrometer cycle was programmed to collect one full MS scan followed by 10 data dependent tandem mass spectrometry MS/MS scans. The MS scans (300–1800 m/z range, 1,000,000 AGC, 150 ms maximum ion time) were collected at a resolution of 70,000 at m/z 200 in profile mode and the MS/MS spectra (2 m/z isolation width, 25% collision energy, 100,000 AGC target, 50 ms maximum ion time) were acquired at a resolution of 17,500 at m/z 200. Dynamic exclusion was set to exclude previous sequenced precursor ions for 30 seconds within a 10-ppm window. Precursor ions with +1, and +6 or higher charge states were excluded from sequencing.

The MaxQuant (Cox et al., 2014; Luber et al., 2010) (v1.5.5.1) LFQ algorithm was used for protein quantitation as previously described (Seyfried et al., 2017). UniProt protein sequences containing both Swiss-Prot and TrEMBL human protein sequences (90,411 target sequences downloaded April 21, 2015), were duplicated into a reverted (decoy) peptide database, searched, and used to control peptide and razor protein false discovery rate (FDR) at 1% within MaxQuant. Methionine oxidation (+15.9949 Da), asparagine and glutamine deamidation (+0.9840 Da), N-terminal acetylation (+42.0106 Da) and cysteine carbamidomethylation (+57.0215 Da) were assigned as fixed modifications. Tryptic peptides with only 2 mis-cleavages were included in each database search. A precursor mass tolerance of ±20 ppm was applied prior to mass accuracy calibration and ±4.5 ppm after internal MaxQuant calibration. Other search settings included a maximum peptide mass of 6,000 Da, a minimum peptide length of 6 residues, 0.05 Da tolerance for high resolution Orbitrap MS/MS scans, or 0.6 Da for low resolution MS/MS scans obtained in the linear ion trap. The FDR for peptide spectral matches, proteins, and site decoy fraction were all set to 1%. Following quantification, missing values were imputed assuming informative missingness such that missing values were replaced with a left Gaussian tail random distribution per parameters previously determined ideal for LFQ based studies (Tyanova et al., 2016). IgG (background) measurements were averaged if either or both AD and control nonspecific IgG replicate measurements for summed intensity were non-missing. Otherwise, either non-missing measurement was used; in the case that both measurements were missing (low background), average noise level imputed value for the two IPs was used for background subtraction. Finally, background subtraction (possibly due to differential specificity of the nonspecific IgG) was not allowed to produce final values below zero (i.e. negative values were set to zero).

### Western blot analysis

For human samples, bound protein complexes were eluted by boiling in Laemmli sample buffer at 98 °C for 5 min. Protein complexes were then resolved on Bolt 4-12% Bis-Tris gels (Thermo Fisher Scientific) followed by transfer to nitrocellulose membrane using iBlot 2 dry blotting system (Thermo Fisher Scientific). Nitrocellulose membranes were incubated with blocking buffer for 30 min followed by overnight incubation using anti-Tau antibody (Tau-E178, Abcam). Membranes were washed with TBST wash buffer (Tris-buffered saline, 0.1% Tween 20) and incubated with fluorophore-conjugated AlexaFluor-680 or AlexaFluor-790 secondary antibodies (anti-mouse or anti-rabbit, 1:15000, Thermo Fisher Scientific) for 1 hr at room temperature. Membranes were washed three times with TBST and scanned using an Odyssey Infrared Imaging System (LI-COR Biosciences).

For *Drosophila*, protein lysates were prepared by homogenizing adult fly heads in 2X Laemmli Sample Buffer (Bio-Rad) (6 μL per head, 10 heads per sample) with a pestle mixer (Argos Technologies), followed by centrifugation at 21,300 x *g* at 4 °C for 15 min. The supernatants were incubated at 95 °C for 5 min before SDS-PAGE analysis. Proteins were separated by 10% Mini-Protein TGX Precast Protein Gels (Bio-Rad), transferred onto PVDF membrane (Millipore), blocked in 5% bovine serum albumin (Sigma) in Tris-buffered saline with 0.1% Tween-20 (Sigma), and immunoblotted using one of the following antibodies: mouse anti-Sm (Y-12, 1:100, Thermo Fisher Scientific), mouse anti-SNF (1:200, gift from Dr. Helen K. Salz), rabbit anti-SNRPD2 (1:250, Novus Biologicals), rabbit anti-U1-70K (1:1000), rabbit anti-Tau (1:5000, Dako), rabbit anti-GFP (1:1000, Invitrogen), rabbit anti-GAPDH (1:5000, GeneTex), mouse anti-Actin (1:1250, Millipore). Membranes were washed with TBST and incubated with HRP-conjugated secondary antibodies (anti-mouse or anti-rabbit 1:10000, Santa Cruz) for 2 hr at room temperature. Membranes were washed with TBST, detected using ECL (PerkinElmer), and scanned using ChemiDoc Imaging system (Bio-Rad).

### Fly stocks and husbandry

Female flies were used for all experiments unless noted otherwise. Flies were raised and aged on molasses-based media at 25°C with ambient light. All experimental genotypes are indicated in figure legends; the following strains were used for this study:

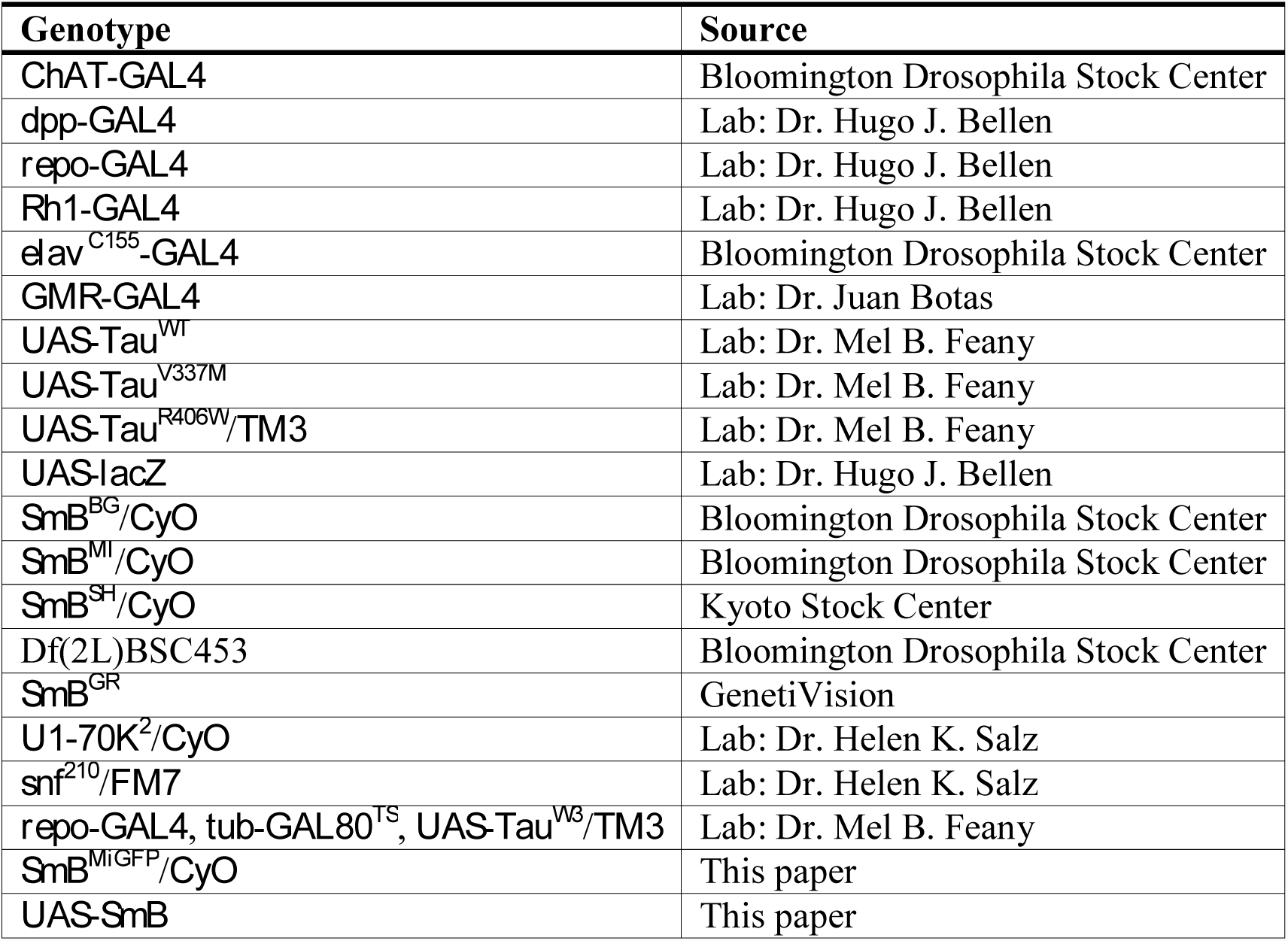

UAS-SmB was generated by cloning *Drosophila* SmB cDNA downstream of the UAS promoter into pUAST-attB vector, which was then inserted using PhiC31 integrase into the attP2 site on 3L chromosome by embryo microinjection. *SmB*^*MG*^ was generated by GFP conversion of *SmB*^*MI*^ as described in Nagarkar-Jaiswal et al., 2015a. Briefly, female flies with the hs-FLP and vasa-phiC31 integrase on the X chromosome, and the FRT flanked GFSTF (EGFP-FlAsH-StrepII-TEVcs-3xFlag) cassette on the third chromosome were mated to *SmB*^*MI*^ males at 18 °C. The resulting embryos were heat shocked at 37 °C for 20 min in 3 consecutive days, and the adult progeny with desired genotype were selected and crossed with flies carrying *CyO* balancer, followed by PCR validation. *Repo*-*GAL4, tub*-*GAL80*^*TS*^, *UAS*-*Tau*^*WT*^/TM3 was maintained at 18 °C to prevent Tau expression and Tau-mediated lethality. To induce glial Tau expression, flies were shifted to 30 °C on the first day of eclosion.

### Sarkosyl extraction

Methods for sarkosyl soluble/insoluble fractionation was adapted from Colodner and Feany, 2010. Briefly, 50 heads from 1- and 20-day-old control (*repo*-*GAL4, tub*-*GAL80*^*TS*^/+ and glial Tau expressing (*repo*-*GAL4, tub*-*GAL80*^*TS*^, *UAS*-*Tau*^*WT*^/+) flies were homogenized in 50 μL of homogenization buffer (15 mM NaCl, 25 mM Tris-HCl at pH 7.4, 1 mM EGTA, 1 mM EDTA, and protease inhibitors) by a pestle mixer (Argos Technologies), followed by brief centrifugation to remove large debris. The homogenate was then centrifuged at 100,000 x *g* at 4 °C for 1 hr. The resulting supernatant was saved as soluble fraction, and the pellet was homogenized in salt/sucrose buffer (10% sucrose, 0.8 M NaCl, 10 mM Tris-HCl at pH 7.4, 1 mM EGTA, and protease inhibitors), followed by centrifugation at 15,000 x *g* at 4 °C for 30 min. Sarkosyl was added to make 1% sarkosyl-containing supernatant, which was then incubated at 37 °C for 1 hr, followed by centrifugation at 100,000 x *g* at 4 °C for 2 hr. The resulting pellet was homogenized in 1% sarkosyl-containing salt/sucrose buffer again, followed by centrifugation at 100,000 x *g* at 4 °C for 2 hr. Lastly, the resulting pellet was homogenized in 15 μL of homogenization buffer and then subjected to western blot analysis.

### Assays for ***Drosophila* survival, climbing, and rough eyes**

For survival analyses, 288∼337 flies per genotype were aged in the vials with less than 30 flies per vial. Surviving flies were counted and transferred to vials with fresh food every other day. For the startle-induced, negative geotaxis assay (climbing), 5-18 flies from each group were placed in individual vial for a total of 5-7 vials for each group; sample/group size variability was due to reduced survival at aged timepoints. Flies from each vial were transferred to an empty vial without anesthetization prior to the assay. Locomotor activity was assessed by counting the number of flies climbing past the 5 cm line within a 5 second interval after gently tapping flies to the bottom of the vial. Photos of external eye appearance were acquired using a Leica EC3 system.

### Histology and immunofluorescence

For histology, *Drosophila* heads were fixed using 8% glutaraldehyde (Electron Microscopy Sciences) at 4 °C for 6 days, followed by paraffin embedding and microtome sectioning, as previously described in Chouhan et al., 2016. Serial 5 mm-thick frontal sections (Leica) were prepared from *Drosophila* heads and mounted on slides, followed by hematoxylin and eosin staining or 3,3’-Diaminobenzidine (DAB) staining to examine brain morphology and cholinergic neurons, respectively. For *Drosophila* brain morphology, we quantified numbers of central brain vacuoles greater than 5 mm in diameter in 5 serial frontal sections at the level of the fan-shape body. From well-oriented frontal sections, Kenyon cells were counted within 6,400 μm^2^ area centered on the mushroom body calyx. For counting cholinergic neurons in *ChAT*>*lacZ* flies, frontal sections were incubated for 25 min. in 10 mM sodium citrate at pH 6.0 at 98 °C for antigen retrieval, followed by washing in phosphate-buffered saline (PBS) containing 0.3% Triton X-100 (PBST) for 5 min. After blocking in 1% normal horse serum in PBS for 20 min, sections were incubated with anti-β-galactosidase antibody (1:250, Promega) for 1 hr at room temperature. Secondary detection was performed using the avidin-biotin-peroxidase complex (ABC) kit (Vector Laboratories) with DAB peroxidase substrate (Vector Laboratories) following the manufacturer’s protocol. The number of immunoreactive neurons in the lamina within 60 μm area was counted as described previously in Wittmann et al., 2001. Light microscopy images for histology were acquired using a Leica DM 6000 B system.

For whole mount immunofluorescence, *Drosophila* brains or imaginal discs were dissected and fixed using 4% formaldehyde in PBS at room temperature for 20 minutes, followed by washing with PBST twice for 10 min each, blocking with 5% normal goat serum in PBST for 1 hour at room temperature. Tissues were incubated with the primary antibodies at 4 °C for 3 days (brains) or 16 hr (third instar larval wing discs), followed by secondary antibody incubation for 2 hours at room temperature, and mounting with Vectashield DAPI-containing mounting medium (Vector Laboratories). The following primary antibody dilutions were used: anti-Sm (Y-12, 1:1000, Thermo Fisher Scientific), anti-SNF (1: 1500, gift from Dr. Helen K. Salz), anti-Tau[pS214] (1:200, Invitrogen). For secondary antibodies, we used Cy3-conjugated goat anti-mouse IgG (1:1000, Jackson ImmunoResearch) or Alexa Fluor 488-conjugated goat anti-rabbit IgG (1:1000, Jackson ImmunoResearch). Confocal microscopy images were acquired with a Leica SP8 confocal system (Leica Microsystems) and using Leica Application Suite X (LAS X) Software.

### TUNEL assay

TUNEL staining was performed using Click-iT Plus TUNEL Assay (Invitrogen). *Drosophila* brains were dissected and fixed by 4% formaldehyde in PBS at room temperature for 20 min, followed by washing with PBST twice for 10 min each, and Protease K treatment for 15 min at room temperature. After a brief wash with water for 5 min, brains were equilibrated in TdT reaction buffer at 37 °C for 10 min, followed by incubation in TdT reaction mixture at 37 °C for 70 min. After being washed with water and PBST containing 5% normal goat serum for 5 min each, brains were incubated with Click-iT Plus TUNEL reaction cocktail at 37 °C for 30 min. Brains were then washed by PBST containing 5% normal goat serum and water for 5 min each before mounted on slides with Vectashield DAPI-containing mounting medium (Vector Laboratories). Images were acquired using Z stack (5 μm per step) and TUNEL-positive signals central brain were counted at the level between ellipsoid body and fan-shaped body.

### Electrophysiology

Methods for electroretinogram recordings were adapted from Chouhan et al., 2016. Briefly, 10 flies from each genotype in 1-, 5-, or 10-day-old were affixed to a glass slide. Flies were kept in dark for at least 1 min prior to stimulation for 1-minute with alternating 2-sec light/dark pulses. Retinal responses were recorded using electrodes placed on the corneal surface and the thorax (reference) and analyzed using LabChart software (ADInstruments).

### *Drosophila* RNA sequencing

For RNA-sequencing (RNA-seq) experimental design, we evaluated each of the following genotypes: (1) *elav*>*Tau*^*WT*^ (*elav-GAL4*/+; +/+; *UAS-Tau*^*WT*^/+ and *elav-GAL4*/*Y*; +/+; *UAS-Tau*^*WT*^/+); (2) *elav*>*Tau*^*R406W*^ (*elav-GAL4*/+; +/+; *UAS-Tau*^*R406W*^/+ and *elav-GAL4*/*Y*; +/+; *UAS-Tau*^*R406W*^/+); (3) *SmB*^*MG*/*MG*^; (4) *elav* control (*elav-GAL4*/+ and *elav-GAL4*/*Y*); and (4) *yw* control (*yw/yw* and *yw/Y*). For *elav>Tau* and *elav* controls, animals were evaluated at 1-, 10-, or 20-days. For *SmB* and *yw* controls, 10-day old animals were evaluated. To avoid possible batch effects, all experimental and control genotypes used for each comparison (*Tau*^*WT*^, *Tau*^*R406W*^, or *SmB*) were sequenced together. Triplicate samples, prepared from independent groups of pooled fly heads, were evaluated for all genotypes and age, except for the *elav* control genotype used for the comparison with *Tau*^*R406W*^, for which duplicate samples were used:

| **<u>Exp</u>** | **<u>Replicates (n)</u>** | **<u>Ctrl</u>** | **<u>Replicates (n)</u>** | **<u>Age (d)</u>** |
| --- | --- | --- | --- | --- |
| *Tau<sup>WT</sup>* | 3 | *elav* | 3 | 1, 10, 20 |
| *Tau<sup>R406W</sup>* | 3 | *elav* | 2 | 1,10, 20 |
| *SmB* | 3 | *yw* | 3 | 10 |

Total RNA was extracted from approximately 100 adult fly heads (for each genotype/age/sample), equally divided between males and females. Frozen fly heads were homogenized in Trizol (Invitrogen), treated with DNaseI (Promega), and total RNA was extracted using the RNeasy Micro Kit (Qiagen). RNA-seq was performed following the Large Insert Strand Specific TruSeq protocol by the Genomics Platform at The Broad Institute. Each library was sequenced with 100 bp pair-end reads. Raw reads were obtained in binary format (BAM) format. Fastq files were extracted from bam files using Picard tools SamToFastq module. Sequencing quality and any adapter contamination were assessed using FastQC v0.10 (Andrews, S. *FastQC: a quality control tool for high throughput sequence data*, 2010). Average phred score for all samples was greater than 35. Average sequence depth for *Tau*^*WT*^, *Tau*^*R406W*^, and *SmB* samples (and respective controls) was 53M, 84M, and 79M read pairs, respectively. Due to reduced coverage, one of the *elav* control samples (10-day-old) from the *Tau*^*WT*^ comparison was re-sequenced and subsequently down sampled to 60M read pairs, using random selection in seqtk (https://github.com/lh3/seqtk). For the *SmB* comparison, one experimental and control sample were each down sampled to 90M read pairs. Based on the *Drosophila* transcriptome size (30M bp), average coverage across all samples is estimated at ∼230X, which was deemed adequate for our subsequent splicing analysis pipeline. Raw reads were aligned to *Drosophila* reference genome dm6 and human reference genome GRCh38 and using STAR 2.5.3a (Dobin et al., 2013). Whole genome FASTA sequence and annotations of respective genome builds were downloaded from UCSC genome browser portal (http://hgdownload.soe.ucsc.edu/downloads.html). Raw genome is indexed by setting --runMode to genomeGenerat in STAR. To reliably align splice junction, the raw reads were aligned to the indexed genome by setting an anchor length (splice junction) of 5 and --outSAMstrandField to intronMotif. Sample-wise alignments were saved as coordinate sorted BAM files. Average mappability of all samples was 96.5%. Gene level counts were obtained as the sum of total reads mapped to respective genes. As expected, based on principal component analysis using gene counts (Yalamanchili et al., 2017)., sample clustering reflected genotype and age (Figure S7).

For evaluation of differential expression of alternative spliceforms we deployed rMATS (Shen et al., 2014). Alignment files (BAM) and reference annotations (GTF) from UCSC genome browser were passed to rMATS. Insert length is computed as average fragment size (400 bp) - (2*read length). rMATS classifies splicing events into 5 categories, skipped exons, retained introns, mutually exclusive exons, alternative 5’ ss, and alternative 3’ ss (Figure S6A). For analysis of intron retention, we developed and applied a new, complementary tool, Differential Expression of Introns (DEIn). In order to minimize false positive calls, either due to sequencing artifact, non-specific / multi-mapping, or incompletely processed mRNA, we required each intron to be fully covered (i.e. every nucleotide appears in at least 1 sequencing read) (See schematic in Figure S6B). Aligned RNA-seq (BAM) files were converted to bigwig format with a bin size of 2 using the bamCoverage module from deeptools (Ramírez et al., 2014). Based on the *Drosophila* reference genome dm6, we excluded all sequences that overlap with annotated exons. Coverage of each intron across all genotypes and time points were computed from respective bigwig files using bwtools (Pohl and Beato, 2014). Only fully covered introns in at least one of the samples were retained. Differential expression was determined using DESeq2 (Love et al., 2014), with fold-change >0. In order to detect cryptic splicing errors, we used the published tool, CrypSplice (Tan et al., 2016). CrypSplice only considers splice junctions that have not previously been annotated in public databases; defining cryptic junction strength as the ratio of junction reads to total 5’ splice site coverage. For each cryptic junction, a strength difference is computed: experimental minus control samples. We focused on junction gains (junction strength difference > 0) with a minimum threshold of at least 10 junction reads. Junctions spanning multiple genes were assigned to the gene of origin. In cases where cryptic junctions originate outside of a gene body, we computed the distance to the nearest gene, and only considered junctions mapping within 500bp of a gene.

For descriptive purposes, CrypSplice was enhanced to assign cryptic junction categories (Figure 6A), including new splice donor (D), new splice acceptor (A), N (new splice donor & new splice acceptor), NDA (exon skipping). All genes with splicing errors identified by DEIn (intron retention) CrypSplice (cryptic junctions) were further annotated based on numbers of introns and alternative transcript spliceforms. We also examined the consensus sequences for error-prone splice donor and acceptor sequences, including splice sites flanking all retained introns (DEIn) and those annotated splice sites “skipped” to generate new splice donor (D) or acceptor (A) detected by CrypSplice (examples highlighted in Figure 6C). Binding strength of splice donor and acceptor sites was determined based on the entropy of respective 5’ (−3 bp exonic to +6 bp intronic) and 3’ (−20 bp intronic to +3 bp exonic) splice site sequences (Yeo and Burge, 2004). Finally, the positions of splicing errors within transcripts were annotated with respect to coding regions, 5’ untranslated regions (UTR), and 3’ UTR.

Selected splicing errors were validated by PCR, using the following oligonucleotide primers:

*Diap1*-F 5’-GTCAAATCTCAACGCAACGG-3’

*Diap1*-R 5’-TAGCTCCTTTGTTTGCCTGACT-3’

*Grik*-F 5’-GAAAATCGGGCAATGGAGCG-3’

*Grik*-R 5’-GCCCTCAAACTGATCGTTGC-3’

*Oli*-F 5’-AAATCGCGGTCGTCTCTGTT-3’

*Oli*-R 5’-GCTGGCCACCAAGCATATTG-3’

### Human (ROSMAP) RNA-seq analysis

For CrypSplice analysis of ROSMAP RNA-seq data, we initially dichotomized based on AD pathologic diagnosis (above). In a complementary approach, we also compared 100 cases and 100 controls selected from the extremes (high/low) of neurofibrillary tangle pathology, using the quantitative composite score for tangle pathologic burden (see also below). Our analyses focused on junction gains (junction strength difference > 0) with a minimum threshold of at least 5 reads, and detected in greater than 50% of cases and less than 50% of controls. In order to evaluate non-recurrent cryptic splicing errors, we developed and applied the Cryptic Load tool. Cryptic Load computes a person-specific cryptic splicing burden, based on average cryptic junction strength,

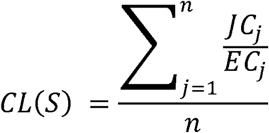

where, *CL(S)* is the Cryptic Load of a sample *S, JC*_*j*_ is the read count of junction *j, EC*_*j*_ is the respective 5’ exon coverage of junction *j* and *n* is the total number of cryptic junctions in sample *S*. Cryptic load was further normalized to respective sequencing depth. For initial alignment and mapping of junctions from the large number of ROSMAP samples we used the cloud formation cluster at Amazon Web Services. We created one EC2 master instance (m3.xlarge) which was used to launch EC2 computing nodes of type (c3.8xlarge).

### Quantitative real-time PCR (qRT-PCR)

Total RNA was prepared from *Drosophila* heads as described above. Following reverse transcription using the SuperScript III First-Strand Synthesis System (Invitrogen), qRT-PCR was performed using iQ SYBR Green Supermix (Bio-Rad) in a CFX96 Touch Real-Time PCR Detection System (Bio-Rad) with standard cycling parameters. Each reaction was performed in triplicate. *RpL32* was used as a control for normalization of each transposon to calculate ΔCt values. The following primers were used:

*RpL32-F* 5’-ATCGGTTACGGATCGAACAA-3’

*RpL32-R* 5’-GACAATCTCCTTGCGCTTCT-3’

*snRNP-U1-70K-F* 5’-GCTGTTCATTGCACGCATCA-3’

*snRNP-U1-70K-R* 5’-TGCTCGTACTCGATGAAGGC-3’

*SmD2-F* 5’-GTACTGGAAAACGTGAAGGAGAT-3’

*SmD2-R* 5’-GAATCGCCTCGCAGGAACA-3’

*SmB-F* 5’-ATGACGATCGGCAAGAACAAC-3’

*SmB-R* 5’-TGTGTTTGTCGAAGGCTTTGA-3’

*Snf-F* 5’-ATGAGAAGAAGGACAAGAAGAAGAA-3’

*Snf-R* 5’-GGCGTAATCTTGAAGCCCTG-3’

*SmD3-F* 5’-GCACAGGCTCGTGGCAGAGGAA-3’

*SmD3-R* 5’-GTTGGTCCTCCCTGCCATG-3’

## QUANTIFICATION AND STATISTICAL ANALYSIS

### Human Tau IP LC-MS/MS analyses

Differentially enriched or depleted proteins in the Tau interactome were determined by calculating t-test p-values (unpaired) and fold change difference (±1.5 fold) for the pair wise comparison of AD versus control, using the aov() and anova() functions in R. The ANOVA p-value (probability > F) calculation provides identical results to t.test() when only 2 groups are compared. Volcano plots were generated with the R ggplot2 package. ‘Spliceosome’, ‘RNA metabolism’, and ‘Translation’, GO-associated genes were downloaded on 11/16/2018 from geneontology.org, and matching gene symbols in the IPs were annotated in Table S2. DAVID v6.8 was used to calculate enrichment p-values for gene ontologies enriched among the Tau-associated proteins differentially increased in AD. The standard human background was used for enrichment analysis as the Tau interactor list is functionally biased. Next, STRING (Snel et al., 2000) was used to map functional interactions among the increased AD Tau interacting proteins assigned to the “Poly(A)-RNA binding”, “Translation” or “Ribonucleoprotein” GO-terms (n = 105 total). Proteins from the “Ribonucleoprotein” cluster in STRING (Figure S1) were clustered hierarchically using the NMF package aheatmap function, with euclidian distance and complete linkage, and summed intensity data was converted to fold standard deviation (SD) from the mean for each row (i.e., Z score) prior to clustering.

### *Drosophila* experimental data analyses

*Drosophila* experimental image data were analyzed using ImageJ (NIH). Sample size for all comparisons is included in each figure legend and in METHOD DETAILS (above). Statistical analyses used either two-tailed, unpaired t-tests or Analysis of Variance (ANOVA), followed by Tukey’s post-hoc test for multiple comparisons, as specified in all Figure legends. Survival assays were analyzed using the Kruskal-Wallis test followed by Dunn’s test for post hoc comparisons. The significance threshold for all analyses was set to p < 0.05. Otherwise, results are noted at “not significant” (ns). Error bars in all analyses represent the standard error of the mean (SEM).

For statistical comparisons of *Drosophila* RNAseq results, the default tests were used as implemented within rMATS (likelihood ratio test), DEIn/DESeq2 (Wald test), and CrypSplice (beta binomial test). Significance threshold was set to adjusted p-value (FDR) < 0.05, based on the Benjamini-Hochberg procedure. For analyses of gene/transcript/junction features, including intron or alternative splice form numbers (Figure 6E) and 5’/3’ splice site binding strength scores (Figure S6D), random sampling was implemented to determine an empirical p-value using python. Random samples were drawn from the background of all *Drosophila* genes/transcripts/exon-intron junctions, respectively, using a gene set size equal to the experimentally determined comparison group (e.g. n=592 genes for Tau-associated cryptic splicing errors). P-value was computed based on 1000 random samples, noting the frequency of mean feature values as or more extreme than the experimental set. Gene sets with Tau-associated splicing errors were evaluated for functional enrichment of biological process (BP) gene ontology terms using DAVID 6.8 (Huang et al., 2009a, 2009b) with a standard *Drosophila melanogaster* background. Tables S13 display all results meeting a significance threshold of p < 0.001, and overlapping genes, n > 5.

### ROSMAP RNAseq analysis

As described above for *Drosophila*, CrypSplice analyses of ROSMAP RNAseq data also used the beta binomial test and a significance threshold of FDR < 0.05. For analyses of CrypticLoad, linear regression was performed using the model,

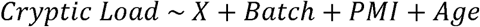

where X=case/control, and additional covariates were included for sequencing batch, postmortem interval (PMI) and age of death (Age). Case / Control status was assigned using 3 separate outcome traits: AD pathologic diagnosis, Braak neurofibrillary tangle stage (high/low), or quantitative neurofibrillary tangle pathologic burden. For tangle burden, we selected equal numbers of cases and controls from each tail of the distribution (high versus low tangle burden) varying total sample size (cases + controls) between 600 and 100 brains (Table 1). For data visualization, cumulative density plots were generated using the ecdf() function and ggplot2 R, stratifying based on AD diagnosis, Braak tangle stage, or neurofibrillary tangle burden.

## DATA AND SOFTWARE AVAILABILITY

RNA-sequencing data for both *Drosophila* and human postmortem brain samples (ROSMAP) are available on the AMP-AD Knowledge Portal (doi:10.7303/syn2580853). RNA-seq from *Tau*^*WT*^, *Tau*^*R406W*^ and *SmB*^*MG*^ flies is available at doi:10.7303/syn7274101. ROSMAP RNA-seq data is deposited at syn3388564. Scripts and analysis pipelines used in this study, including CrypSplice, CrypticLoad, and DEIn, are available at: www.liuzlab.org/Scripts_Tau-SmB.zip.

## SUPPLEMENTAL INFORMATION

Supplemental information includes 6 Supplemental Figures and 16 Supplemental Tables. Supplemental File 1 is comprised of 7 Supplemental tables provided as excel worksheets. Supplemental File 2 includes the differential alternative splicing dataset.

