## Supplemental Information for "Tau-mediated Disruption of the Spliceosome Triggers Cryptic RNA-splicing and Neurodegeneration in Alzheimer’s Disease"

### SUPPLEMENTAL FIGURES

**Figure S1**

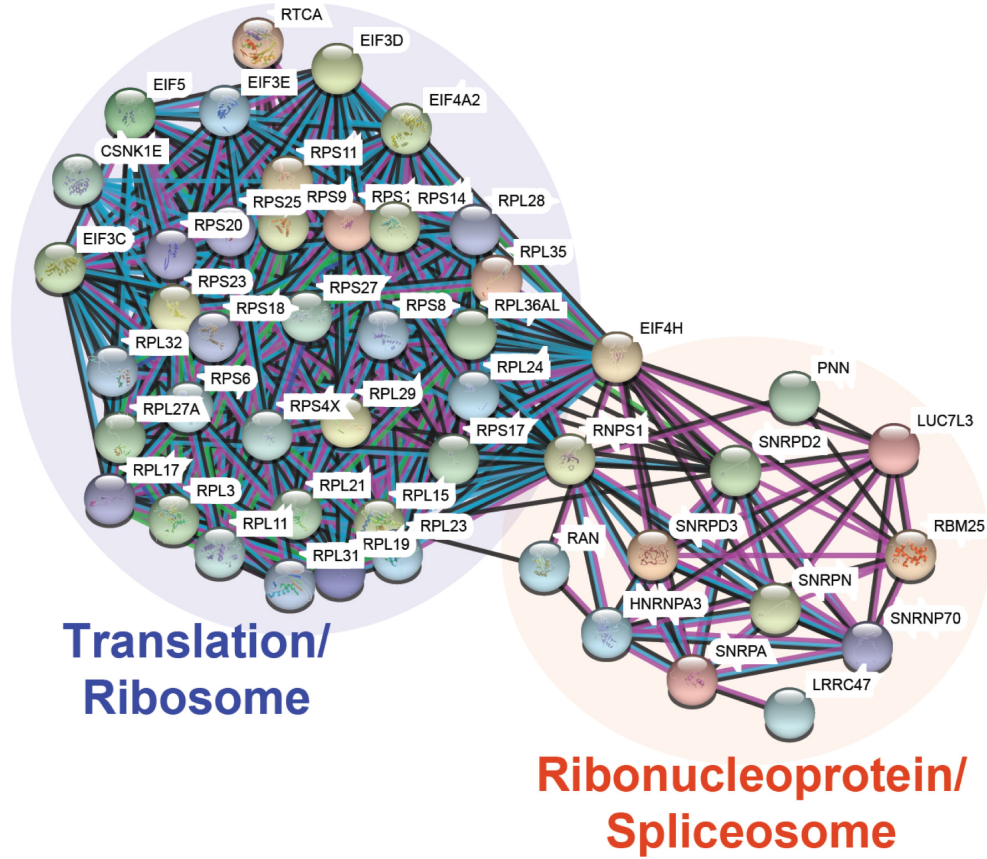

**Supplementary Figure 1. A network of Tau-associated ribonucleoproteins in AD brains (Related to Figure 1).** Based on 3 strongly-enriched Gene Ontology terms (poly-A RNA binding proteins, $p = 4.5 \times 10^{-24}$; ribonucleoprotein, $p = 1.3 \times 10^{-14}$; and translation, $p = 3.1 \times 10^{-14}$), 105 proteins (fold-change $> 1.5$; $FDR < 0.05$) showing increased association with Tau in AD postmortem brain were further analyzed using STRING. The resulting protein-protein interaction network highlights 2 subnetworks: a first cluster with roles in translation (Purple) and a second group of ribonucleoproteins (Orange), including several core spliceosome components.

Figure S2

**A**

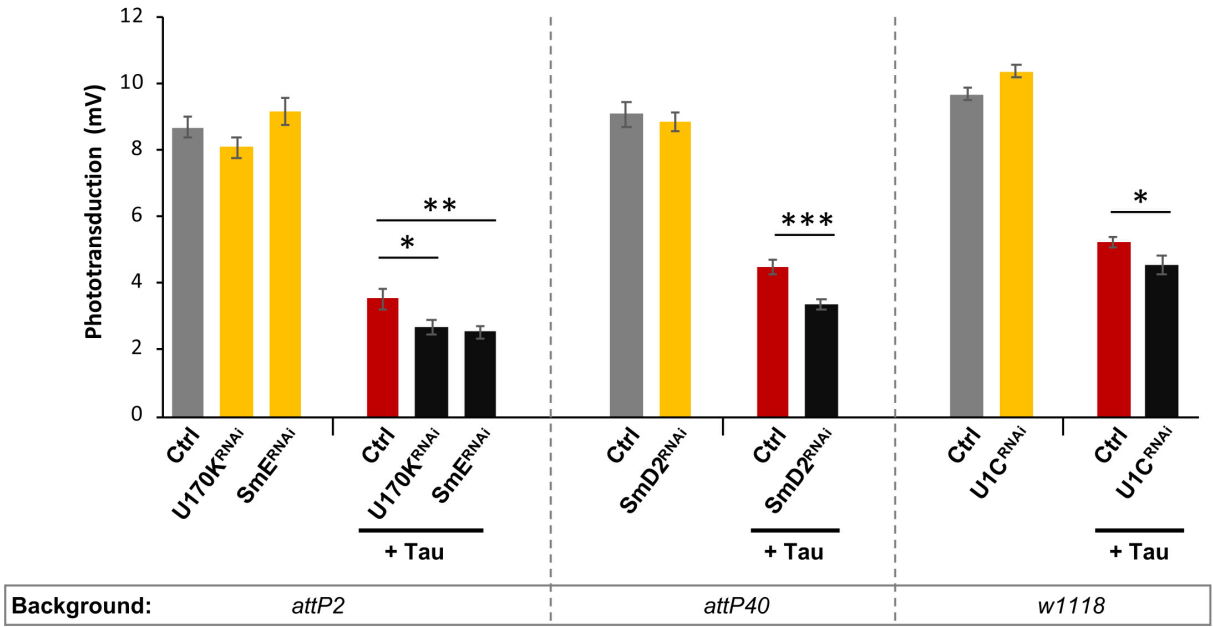

**B**

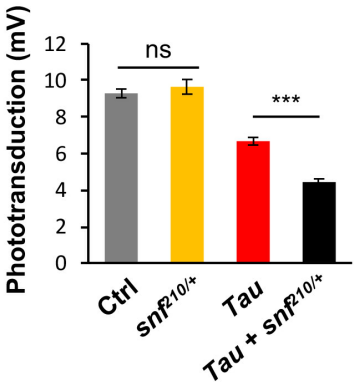

**C**

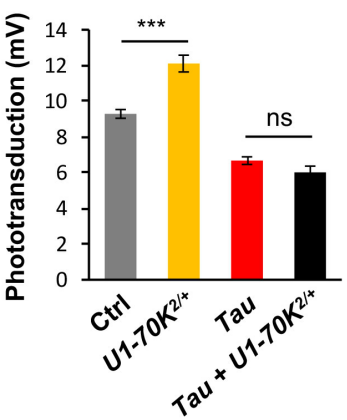

**D**

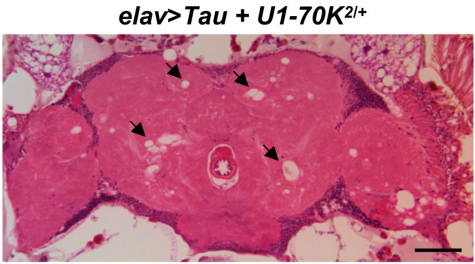

**E**

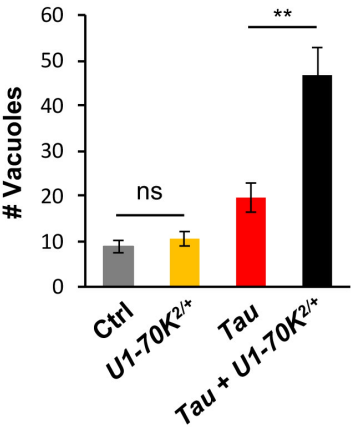

**Supplementary Figure 2. Additional characterization of genetic interactions between Tau and snRNP components (Related to Figure 2).**

(B) RNA-interference (RNAi) targeting multiple spliceosome factors enhances Tau-mediated retinal toxicity, based on electroretinograms (ERGs) in 5-day-old animals. Expression of both Tau and RNAi were directed to adult photoreceptors using *Rh1-GAL4*. In order to control for genetic background, Tau and ctrl strains were outcrossed to (1) *P{y[+t7.7]=CaryP}attP2*; (2) *attP40* for *P{y[+t7.7]=CaryP}attP40*; or (3) *w1118*. RNAi-mediated knockdown of *U1-70K* (*HMS00274*), *SmE* (*HMS00074*), *SmD2* (*HMC03839*), and *U1C* (*v22132*) enhanced Tau-induced retinal toxicity (red, *Rh1-GAL4/+; UAS-Tau<sup>WT</sup>/+*; black, *Rh1-GAL4/+; UAS-Tau<sup>WT</sup>/UAS-RNAi* or *Rh1-GAL4/+; UAS-RNAi/+; UAS-Tau<sup>WT</sup>/+*). Compared with controls (Gray: *Rh1-GAL4/+*), knockdown of each spliceosome factor independent of Tau does not show toxicity on its own (Yellow: *Rh1-GAL4/+; +; UAS-RNAi/+* or *Rh1-GAL4/+; UAS-RNAi/+; +*). At least 10 animals were examined for each genotype.

(C) *Snf* exhibits dose-sensitive enhancement of Tau-induced retinal dysfunction, based on ERGs performed in 10-day-old animals. Compared to controls (Gray: *Rh1-GAL4/+* and Yellow: *snf<sup>210</sup>/+*; *Rh1-GAL4/+*), Tau expression (Red: *Rh1-GAL4/+; UAS-Tau<sup>WT</sup>/+*) causes a reduction in photoreceptor depolarization amplitude, and this phenotype is further enhanced in flies heterozygous for the *snf<sup>210</sup>* null allele (Black: *snf<sup>210</sup>/+*; *Rh1-GAL4/+; UAS-Tau<sup>WT</sup>/+*). At least 10 animals were examined per genotype.

(D) *U1-70K* did not show similar enhancement of Tau-induced retinal dysfunction based on ERGs in 10-day-old animals. The following genotypes were examined: (1) Gray: *Rh1-GAL4/+*; (2) Yellow: *Rh1-GAL4/U1-70K<sup>2</sup>*; (3) Red: *Rh1-GAL4/+; UAS-Tau<sup>WT</sup>/+*; and (4) Black: *Rh1-GAL4/U1-70K<sup>2</sup>; UAS-Tau<sup>WT</sup>/+*. At least 10 animals were examined per genotype.

(E) *U1-70K* exhibits dose-sensitive enhancement of Tau-induced neurodegeneration in the adult brain. Tau was expressed pan-neuronally using the *elav-GAL4* driver. Frontal sections were prepared from 10-day-old animals and stained with hematoxylin and eosin to reveal neurodegenerative changes. Tau-induced progressive neuropil vacuolization (arrows) was enhanced in flies heterozygous for the *U1-70K<sup>2</sup>* null allele (*elav-GAL4/+; U1-70K<sup>2</sup>/+*; *UAS-Tau<sup>R406W</sup>/+*). Scale bar: 50 μm.

(F) Quantification of Tau-induced vacuole formation in adult central brains (Red: *Rh1-GAL4/+; UAS-Tau<sup>WT</sup>/+*), compared with controls (Gray: *Rh1-GAL4/+* and Yellow: *Rh1-GAL4/U1-70K<sup>2</sup>*). This phenotype was enhanced in flies heterozygous for the *U1-70K<sup>2</sup>* null allele (Black: *Rh1-GAL4/U1-70K<sup>2</sup>; UAS-Tau<sup>WT</sup>/+*), based on the examination of central brain sections from at least 5-8 brains per group. For statistical analyses, one-way ANOVA followed by Tukey's test for post hoc comparisons were performed. The unpaired t-test was used for the single comparisons in (A, *SmD2* and *U1C*). All error bars denote mean ± SEM. \*, $p < 0.05$; \*\*, $p < 0.01$; \*\*\*, $p < 0.001$; ns, not significant.

Figure S3

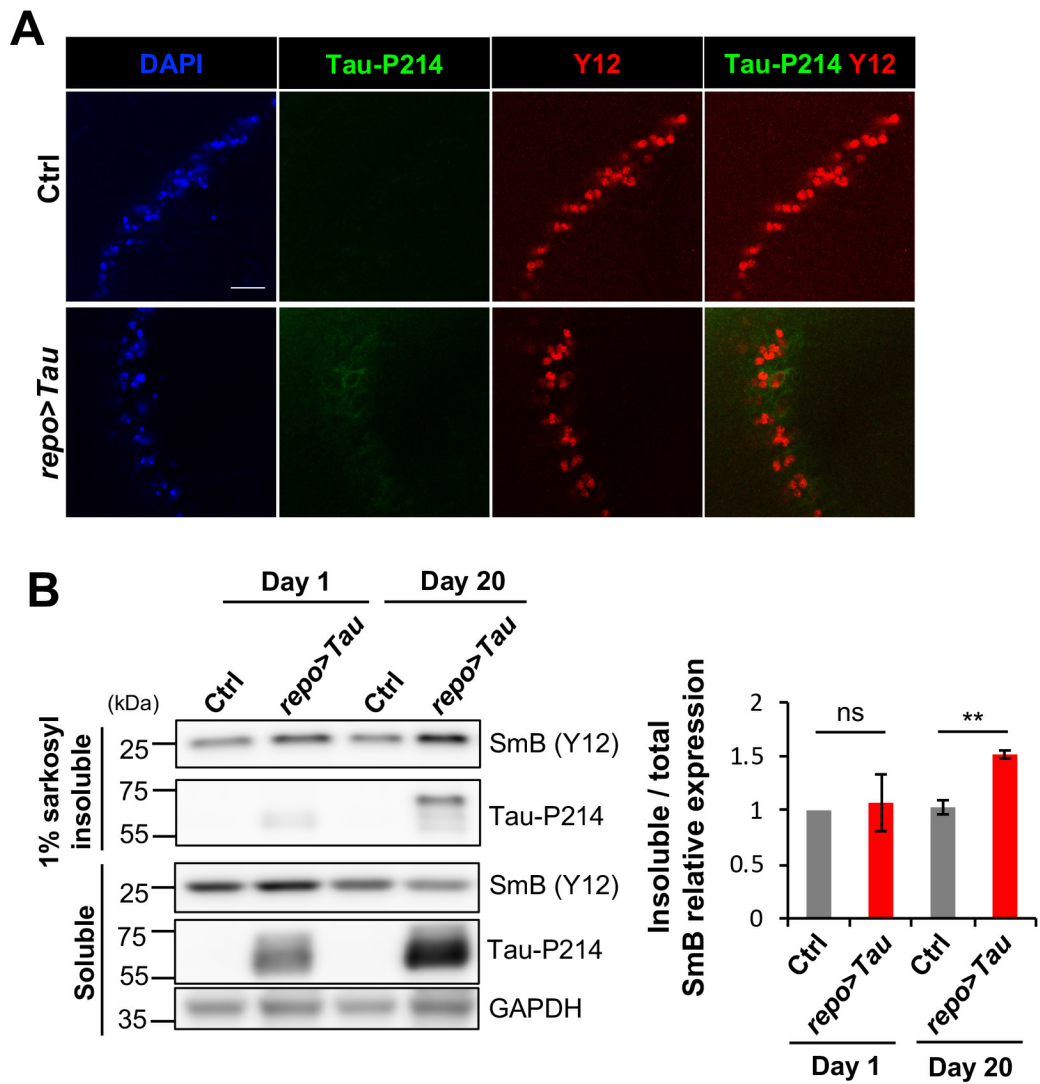

**Supplementary Figure 3. Additional characterization of Sm proteins in Tau transgenic *Drosophila* (Related Figure 3).**

(A) In contrast to aged animals (Figure 3E), 1-day-old flies with glial Tau expression (*repo-GAL4*, *UAS-Tau<sup>WT</sup>*, *tub-GAL80<sup>TS/+</sup>*) do not show develop tangle-like, phospho-Tau aggregates. Representative images from Tau or Control (Ctrl: *repo-GAL4*, *tub-GAL80<sup>TS/+</sup>*) brains were stained for SmB/D3 (Y12, Red) and phospho-Tau (anti-Tau-P214, Green). Nuclei are highlighted with DAPI (Blue). Scale bars: 10 μm.

(B) SmB and Tau protein are enriched in insoluble fractions from heads of 20-day-old flies with glial Tau expression. Western blots were prepared from either soluble or 1% sarkosyl-insoluble fractions from control (Ctrl) or *repo>Tau<sup>WT</sup>* (Tau) fly head soluble and 1% sarkosyl fractions, and probed for SmB (Y12), phospho-Tau (anti-Tau-P214), or GAPDH (loading control). For statistical analyses, unpaired t-tests were performed, based on 3 replicate experiments. All error bars denote mean ± SEM.

Figure S4

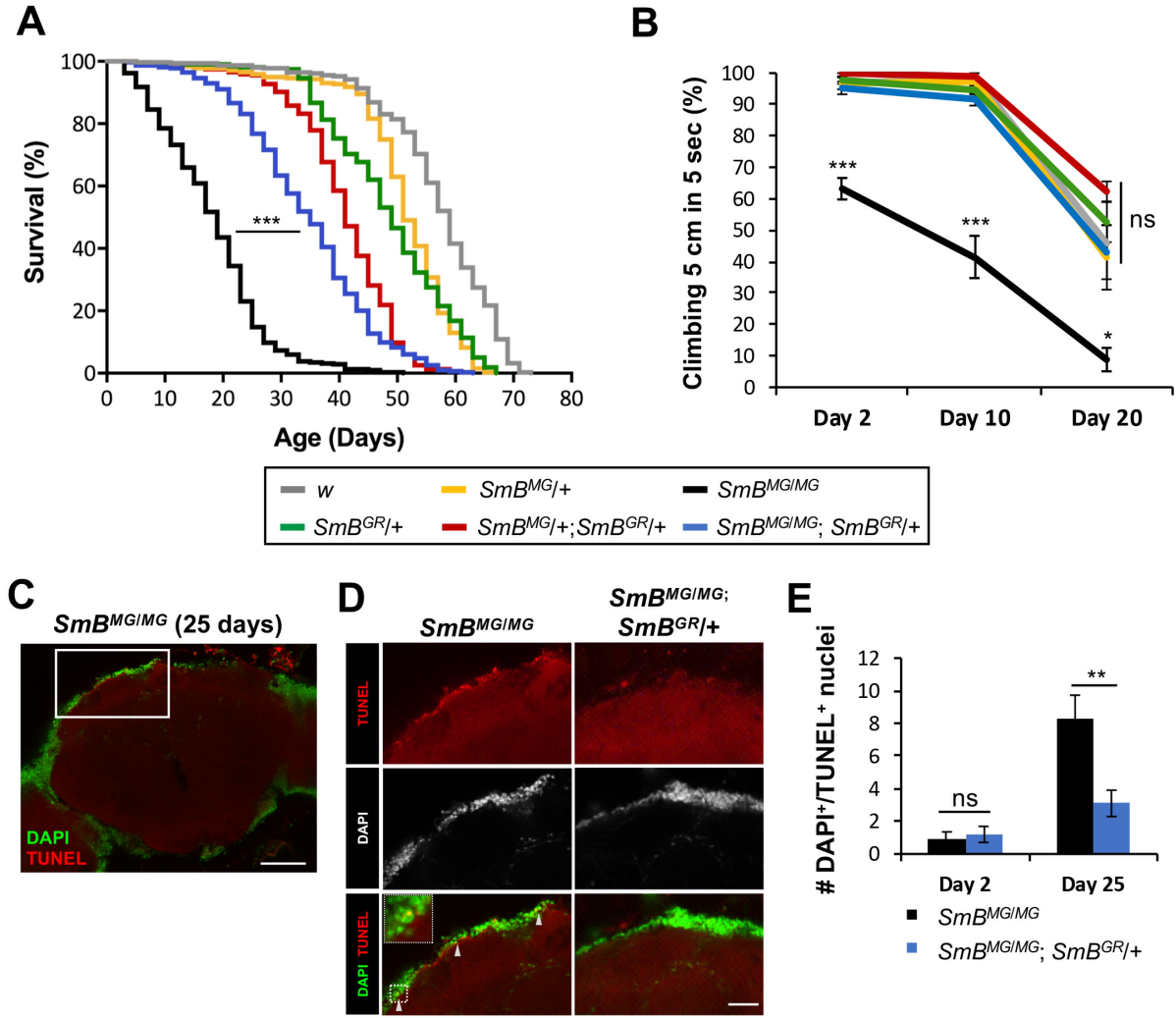

**Supplementary Figure 4. Additional characterization of *SmB<sup>MG</sup>* flies (Related to Figures 4 & 5).**

(A-B) Survival (A) and locomotor (B) assays from Figure 4 are shown including additional control genotypes (Yellow: *SmB<sup>MG/+</sup>* and Red: *SmB<sup>MG/+</sup>; SmB<sup>GR/+</sup>*). Survival of at least 313 adult flies were examined for each genotype. At least 5 replicate experiments were performed, each consisting of 5-18 flies, evaluating the proportion of flies climbing at least 5 cm in 5 sec.

(C-E) Loss of function in *SmB* causes age-dependent neuronal apoptosis in adult brains. (C) Lower power image (scale bar: 50 μm), highlighting region of interest in adult fly brain for comparisons shown in (D).

(D) Representative images of adult brain cortex stained for TUNEL (Red) and DAPI (Gray) from 25-day-old *SmB<sup>MG/MG</sup>* or control (*SmB<sup>MG/MG</sup>; SmB<sup>GR/+</sup>*) flies. Scale bar: 20 μm. Inset shows magnified image to highlight a representative apoptotic nucleus co-staining for TUNEL and DAPI (arrows).

(E) Quantitation of TUNEL/DAPI double-labelled apoptotic nuclei from 2- and 25-day-old animals. At least 6 animals were examined for each genotype and time point.

For statistical analyses, Kruskal-Wallis test (A) followed by Dunn's test for post hoc comparisons, one-way ANOVA (B) followed by Tukey's test for post hoc comparisons, or unpaired t-test (E) were performed. All error bars denote mean ± SEM. \*, $p < 0.05$; \*\*, $p < 0.01$; \*\*\*, $p < 0.001$; ns, not significant.

Figure S5

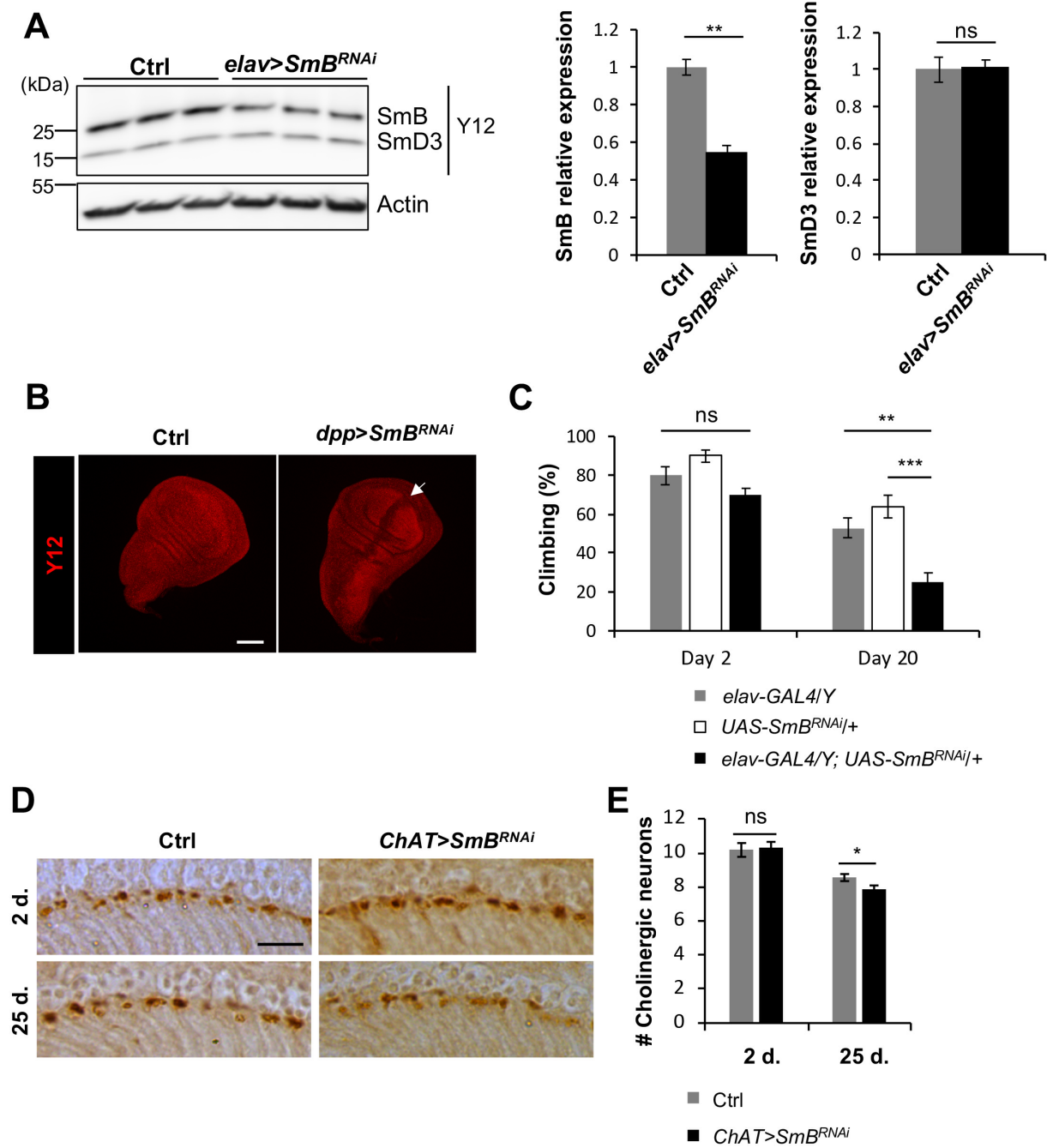

**Supplementary Figure 5. RNAi-mediated knockdown of *SmB* phenocopies *SmB<sup>MiGFP</sup>* (Related to Figures 4 & 5).**

(A-B) RNA interference targeting *SmB* causes reduced SmB protein levels. (A) Western blots were prepared from adult heads of *elav>SmB<sup>RNAi</sup>* (*elav-GAL4/+; +/-; UAS-SmB<sup>HM05097/+</sup>*) and control animals (*elav-GAL4/+*). Blots were probed with the anti-Sm (Y12) antibody, highlighting both SmB and SmD3 proteins or anti-Actin as a loading control. Relative expression of SmB and SmD3, normalized to Actin, was quantified based on analysis of 3 replicate experiments. (B) RNAi knockdown of SmB was also confirmed by anti-SmB staining (Y12) of larval 3<sup>rd</sup> instar wing imaginal discs from *decapentaplegic* (*dpp>SmB<sup>RNAi</sup>* (*dpp-GAL4/UAS-SmB<sup>HM05097</sup>*) and control (*dpp-GAL4/+*) animals. The *dpp-GAL4* driver line expresses in a central stripe of the imaginal disc (arrow), denoted by the reduced Y12 staining. Scale bar: 100 μm.

(C) Compared to controls (*elav-GAL4/Y* and *UAS-SmB<sup>HM05097/+</sup>*), *elav>SmB<sup>RNAi</sup>* flies (*elav-GAL4/Y; +/-; UAS-SmB<sup>HM05097/+</sup>*) exhibit progress locomotor impairment. Quantification based on at least 6 replicate experiments. Data are presented as mean ± SEM.

(D-E) RNAi knockdown of *SmB* causes progressive loss of cholinergic neurons in the lamina. (D) Cholinergic neurons are labeled by DAB staining using the *ChAT-GAL4* reporter line driving expression of β-galactosidase. Compared with controls (*ChAT-GAL4, UAS-lacZ/+*), *Chat>SmB<sup>RNAi</sup>* (*Chat-GAL4, UAS-lacZ/+; UAS-SmB<sup>HM05097/+</sup>*) flies exhibit loss of cholinergic neurons between 2- and 25-days (d.) of age. Scale bar: 10 μm. (E) Quantification of experiment shown in F, based on β-gal-positive neuronal counts from at least 11 animals for each genotype and timepoint.

For statistical analyses, unpaired t-test (A, E) or one-way-ANOVA (C) followed by Tukey's test for post hoc comparisons were performed. \*, $p < 0.05$; \*\*, $p < 0.01$; \*\*\*, $p < 0.001$; ns: not significant. All error bars denote mean ± SEM.

Figure S6

**A**

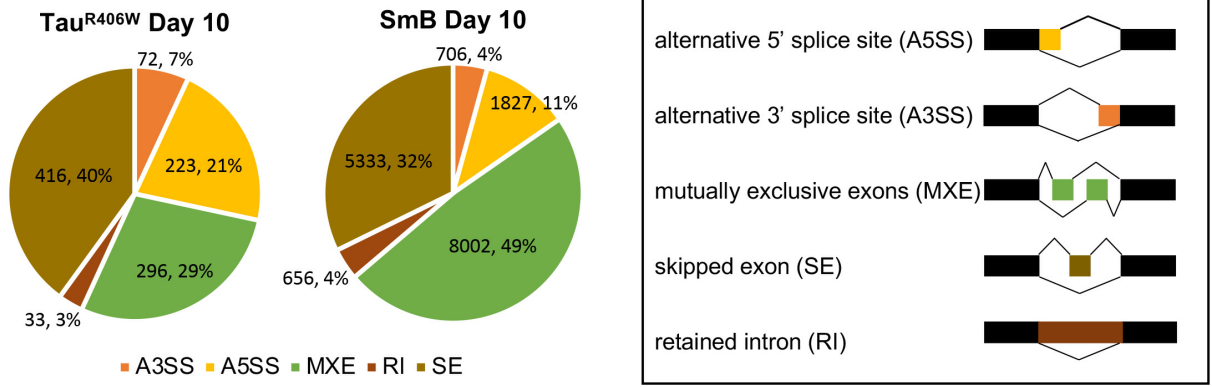

**B**

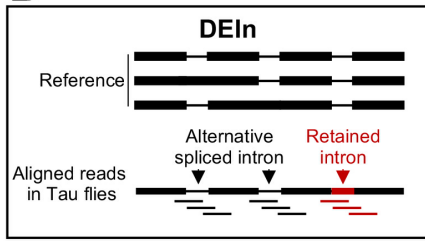

**C**

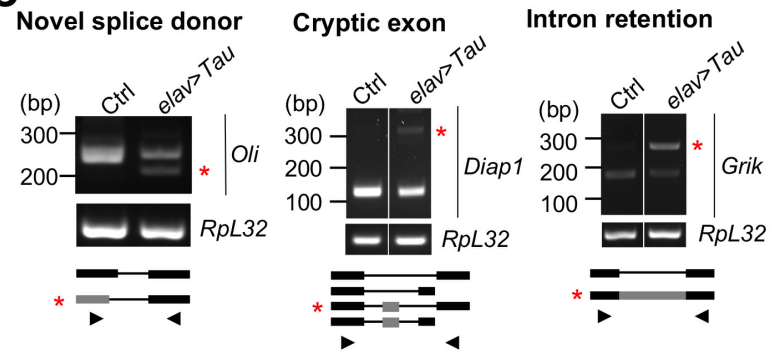

**D**

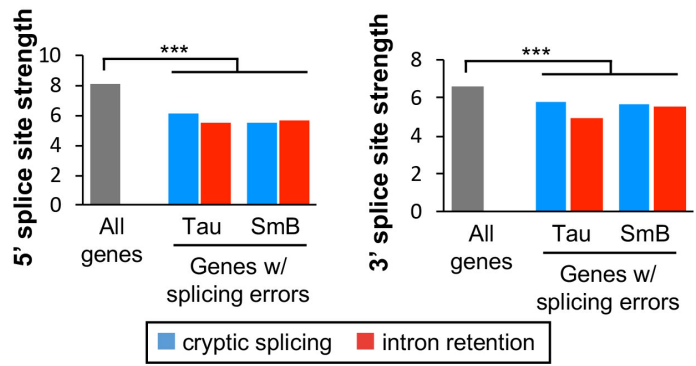

**E**

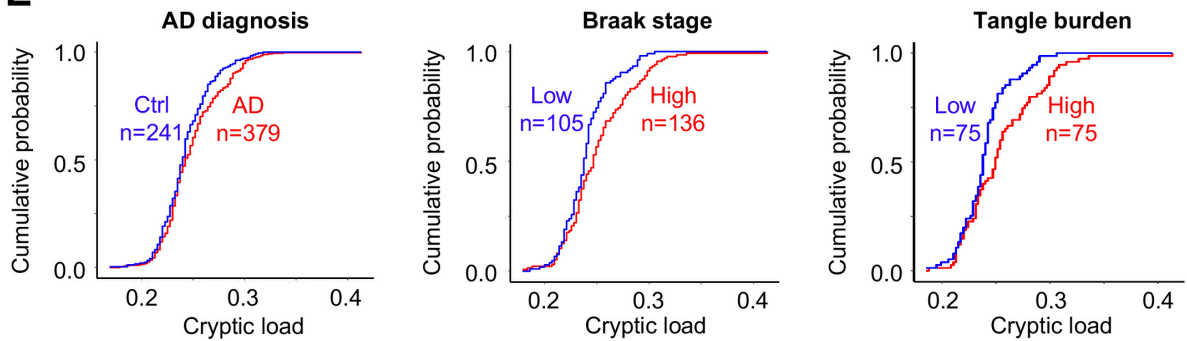

**Supplementary Figure 6. Additional characterization of RNA splicing changes in *Drosophila* and human brains (Related to Figure 6 and Table 1).**

(A) Pie charts highlight the number and frequency of distinct alternative splicing events (schematic at right) differentially expressed in 10-day-old Tau<sup>R406W</sup> (*elav-GAL4/+; +/+; UAS-Tau<sup>R406W</sup>/+*) or *SmB<sup>MG/MG</sup>* flies detected using rMATS.

(B) Schematic showing the stringent definition of intron retention implemented in the DEIn tool. Intron retention events (Red) are counted only for introns that do not appear in the transcriptome reference as expressed sequences, thereby differentiating these events from alternatively spliced exons. DEIn additionally requires that RNA-seq reads span the entire intronic sequence.

(C) Validation of representative splicing errors from Figure 6C. Reverse transcription-polymerase chain reaction (RT-PCR) was performed on total mRNA prepared from heads of 20-day-old Tau (*elav-GAL4/+; UAS-Tau<sup>R406W</sup>/+*) or control animals (*elav-GAL4/+*) (Tau) flies. The aberrant PCR products representing transcripts with splicing errors are indicated (red asterisk). RT-PCR for *RpL32* is shown as a control.

(D) Splicing errors in Tau transgenic or *SmB* flies occur preferentially at splice donor/acceptor sites that diverge from the U1 or U2 consensus binding sites, respectively. Average estimated binding strength score, based on MaxEntScan, is shown for all *Drosophila* annotated 5' splice sites (Left) or 3' splice site strength (Right), along with that associated with splicing errors (blue: cryptic splicing, red: intron retention) in *elav>Tau<sup>R406W</sup>* or *SmB<sup>MG</sup>* flies. The analysis was limited to those cryptic junctions (5': n<sub>Tau</sub>=286 and n<sub>SmB</sub>=225; 3': n<sub>Tau</sub>=172 and n<sub>SmB</sub>=190) for which annotated, apparently skipped, donor or acceptor sequences could be inferred unambiguously (see Methods). For intron retention events, all flanking splice junctions were considered (n<sub>Tau</sub>=2767 and n<sub>SmB</sub>=3953). For statistical analysis, an empirical p-value was computed for each comparison based on random sampling (1000 iterations) from all *Drosophila* splice donor/acceptor sequences, using a sample size equal to the experimentally-defined gene sets with splicing errors in *Tau* or *SmB* animals. \*\*\*, p < 0.001. See also Figure 6F and Tables S11 & S12.

(E) Cumulative probability plots depict increased cryptic load in association with AD pathologic diagnosis (Left), Braak neurofibrillary tangle stage (Middle), or tangle pathologic burden (Right). The plot is consistently shifted to the right in subjects with greater AD pathologic burden (Red), and the greatest separation is observed for the comparison based on the extremes of the tangle burden distribution.

**Supplementary Figure 7. Principal component analyses of *Drosophila* RNA-seq data.**

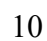

### SUPPLEMENTAL TABLES

**Table S1. Clinical and demographic features of subjects for human brain proteomics (Related to Figure 1).**

| ID | PMI | Onset age | Age at death | Braak score | Sex |
| --- | --- | --- | --- | --- | --- |
| CTL1 | 10 | NA | 57 | 2 | M |
| CTL2 | 6 | NA | 69 | 2 | M |
| CTL3 | 11.5 | NA | 78 | 2 | F |
| CTL4 | 6 | NA | 75 | 1 | F |
| AD1 | 8 | 70 | 78 | 6 | F |
| AD2 | 6 | 69 | 72 | 6 | M |
| AD3 | 7 | 59 | 72 | 6 | F |
| AD4 | 6.5 | 56 | 67 | 6 | M |

**Table S2. Tau associated proteins in AD brains (Related to Figure 1).** See Supplemental File 1 (excel spreadsheet).

**Table S3. Gene ontology enrichment analysis of Tau associated proteins in AD (Related to Figure 1).** See Supplemental File 1 (excel spreadsheet).

**Table S4. RNA-interference strains evaluated for interactions with Tau neurotoxicity (Related to Figure 2).** See Supplemental File 1 (excel spreadsheet).

**Table S5. *SmB<sup>MG</sup>* complementation tests (Related to Figure 4).**

|  | *SmB<sup>MG</sup>* | *SmB<sup>MI07584</sup>* | *SmB<sup>BG02775</sup>* | *SmB<sup>SH0509</sup>* | *Df(2L)BSC453* |
| --- | --- | --- | --- | --- | --- |
| **Molecular Lesion** | Intronic Insertion | Intronic Insertion | Exonic Insertion | Exonic Insertion | Complete deletion |
| **Homozygote viability** | Viable | Lethal | Lethal | Lethal | Lethal |
| ***SmB<sup>MG</sup>* viability** | 19/285 | 0/180 | 0/366 | 0/465 | 0/522 |

**Table S6. Alternative splicing events detected by rMATS (Related to Figure 6).**

|  | **Tau<sup>WT</sup>** |  |  | **Tau<sup>R406W</sup>** |  |  | **SmB** |
| --- | --- | --- | --- | --- | --- | --- | --- |
|  | **Day 1** | **Day 10** | **Day 20** | **Day 1** | **Day 10** | **Day 20** | **Day 10** |
| **Alternative 5' splice site** | 117 | 125 | 104 | 237 | 223 | 264 | 1827 |
| **Alternative 3' splice site** | 80 | 45 | 69 | 97 | 72 | 98 | 706 |
| **Mutually exclusive exons** | 360 | 334 | 476 | 707 | 296 | 293 | 8002 |
| **Skipped exons** | 231 | 139 | 248 | 473 | 416 | 456 | 5333 |
| **Intron retention** | 44 | 40 | 48 | 45 | 33 | 46 | 556 |
| **Total** | 832 | 683 | 945 | 1559 | 1040 | 1157 | 16424 |

**Table S7. Cryptic splicing error counts detected by CrypSplice (Related to Figure 6).**

|  | Tau<sup>WT</sup> |  |  | Tau<sup>R406W</sup> |  |  | SmB |
| --- | --- | --- | --- | --- | --- | --- | --- |
|  | Day 1 | Day 10 | Day 20 | Day 1 | Day 10 | Day 20 | Day 10 |
| **Junctions (n)** | 74 | 103 | 202 | 202 | 352 | 437 | 855 |
| **Genes (n)** | 68 | 91 | 162 | 166 | 295 | 362 | 608 |

**Table S8. Cryptic junctions identified by CrypSplice (Related to Figure 6). See Supplemental File 1 (excel spreadsheet).**

**Table S9. Intron retention counts detected by DEIn (Related to Figure 6).**

|  | Tau<sup>WT</sup> |  |  | Tau<sup>R406W</sup> |  |  | SmB |
| --- | --- | --- | --- | --- | --- | --- | --- |
|  | Day 1 | Day 10 | Day 20 | Day 1 | Day 10 | Day 20 | Day 10 |
| **Introns (n)** | 120 | 118 | 275 | 1123 | 631 | 1138 | 3953 |
| **Genes (n)** | 89 | 97 | 206 | 855 | 522 | 840 | 2446 |

**Table S10. Intron retention events identified by DEIn (Related to Figure 6). See Supplemental File 1 (excel spreadsheet).**

**Table S11. Tau(R406W) cryptic junction features (Related to Figure 6). See Supplemental File 1 (excel spreadsheet).**

**Table S12. Tau(R406W) intron retention junction features (Related to Figure 6). See Supplemental File 1 (excel spreadsheet).**

**Table S13. Gene ontology enrichment analysis of genes affected by splicing errors (Related to Figure 6). See Supplemental File 1 (excel spreadsheet).**

**Table S14. ROSMAP autopsy cohort clinical and demographic characteristics**

|  |  |
| --- | --- |
| **Subjects (n)** | 620 |
| **Age at death, mean yr (SD)** | 88.7 (6.7) |
| **Male, n** | 224 (36.1%) |
| **Education, mean yr (SD)** | 16.3 (3.6) |
| **Clinical AD, n** | 249 (40.2%) |
| **Postmortem interval, mean hrs (SD)** | 7.3 (4.8) |
| **Pathologic AD, n** | 379 (61.1%) |
| **Braak Score, n** |  |
| **0** | 7 (1.1%) |
| **I** | 48 (7.7%) |
| **II** | 50 (8.1%) |
| **III** | 173 (27.9%) |
| **IV** | 206 (33.2%) |
| **V** | 129 (20.8%) |
| **VI** | 7 (1.1%) |

Clinical and pathologic diagnoses of AD were based on NINCDS-ADRDA and NIA-Reagan criteria, respectively.

**Table S15. AD-associated cryptic junctions detected by CrypSplice in ROSMAP (Related to Table 1).** See Supplemental File 1 (excel spreadsheet).

**Table S16. Tangle-associated cryptic junctions detected by CrypSplice in ROSMAP (Related to Table 1).** See Supplemental File 1 (excel spreadsheet).
